# Dietary specialization drives adaptation, convergence, and integration across the cranial and appendicular skeleton in Waterfowl (Anseriformes)

**DOI:** 10.1101/2024.09.21.614171

**Authors:** Ray Chatterji, Tracy A. Heath, Helen F. James, Courtney Hofman, Michael D. Sorenson, Janet C. Buckner

## Abstract

Convergence provides strong evidence for adaptive evolution as it reflects shared adaptive responses to the same selection pressures. The waterfowl (order Anseriformes) are an ideal group in which to study convergent evolution as they have repeatedly evolved morphotypes putatively correlated with diet (i.e., dabbler, grazer, diver). Here, we construct the most robust evolutionary hypothesis to date for waterfowl and reveal widespread morphological convergence across the order. We quantified the shape of the skull and hindlimb elements (femur, tibiotarsus, and tarsometatarsus) of 118 species of extant waterfowl using geometric morphometrics. Multivariate generalized evolutionary models provide strong support for a relationship between dietary ecology and skull shape, and evidence for convergent evolution across lineages that share dietary niches. Foraging behavior better explained the evolution of hindlimb shape, but diet still contributed significantly. We also find preliminary evidence for integration across all three measured hindlimb elements with each other and with the skull. We demonstrate that dietary ecology drives morphological convergence within waterfowl, that this convergent evolution involves multiple integrated skeletal structures, and that morphological changes are associated with shifts in the rate of phenotypic evolution.

---

Convergent evolution is a fascinating evolutionary phenomenon characterized by disparate lineages independently evolving analogous traits, and it remains a key concept for explaining patterns of trait evolution. Evidence of convergence provides some of the strongest support for adaptive evolution as it typically reflects adaptive responses to similar selection pressures. Consequently, convergent processes have been a focal point for evolutionary studies in a wide range of taxa (Wake 1991; Rosenblum et al. 2014; Stayton et al. 2015; Friedman et al. 2016; Baeckens et al. 2019). Much of this focus has been on morphological traits as the most observable manifestation of convergence (McCracken et al. 1999; Stayton et al. 2015; Sherratt et al. 2019; Evers and Benson 2019).

Prevailing theory suggests that convergent phenotypes arise due to adaptation to similar selective pressures, though phenotypic convergence can arise for non-selective reasons (Jaekel and Wake 2007; Stayton 2008; Losos 2011; Wake et al. 2011). Additionally, similar environmental selective pressures can also lead to distinct phenotypes (Felice et al. 2019, Law et al. 2022), and the extent of convergence may be limited by the evolutionary context and constraints of disparate lineages (Stayton 2006, Hansen et al. 2008). For these reasons, carefully designed quantitative research that explicitly tests for the action of potential explanatory mechanisms in trait evolution remains necessary.

Birds display a wide array of trait variation across the 10,000+ described species and, in many cases, lineages appear to have converged on major dietary types (e.g., granivory, frugivory, insectivory, etc.). As such, one might expect that dietary niche might drive convergent patterns of morphological variation as dietary specialization has been demonstrated in several bird radiations (Gibbs and Grant 1987, Lerner et al. 2011, Tokita et al 2017). However, there remains little evidence of convergent evolution in skull shape due to diet at broad taxonomic scales (Navalon et al. 2019; Felice et al. 2019). However, research focused on the diversification dynamics of smaller groups, such as vultures (Steinfield et al. 2023) and Hawaiian honeycreepers (Reding et al. 2008), have provided evidence of convergence. Thus, taxonomic scope may be an important consideration in designing studies aimed at testing for convergent signals across clades.

The waterfowl (order Anseriformes) represent a highly successful, globally widespread order of birds distinguished by their ecology and morphology (e.g., spatulate bills, webbed feet, filter feeding) tied to fresh, brackish, and marine wetland and coastal environments. Waterfowl occupy a variety of dietary niches, including terrestrial grazers, shallow water dabblers, piscivores, and coastal invertivores (Johnsgard 2010). Notably, independent lineages of these birds have evolved to occupy these niches multiple times throughout their evolutionary history, as revealed by early molecular systematic studies of the group (Sraml et al. 1996, McCracken et al. 1999, Donne-Grousse et al. 2002). Thus, waterfowl are an ideal model group to test diet as a driver of morphological convergence as there are replicate cases of the same dietary conditions (Faith 1989; McCracken et al. 1999; Sorenson et al. 1999; Olsen 2017). While previous examinations of the adaptive morphology of living waterfowl have focused on specific structures within specific niches (e.g., skulls in grazing morphs –Olsen 2017; hindlimbs in diving morphs – McCracken 1999; Mendoza and Gomez 2022), they have provided some evidence of convergent evolutionary processes. Further descriptive studies of the waterfowl fossil record have also pointed to convergence to explain analogous traits, with the Hawaiian moa-nalo species converging with ‘geese’ (James and Burney 1997; Sorenson et al. 1999) and the North American *Chendytes lawi* converging with sea ducks (Buckner et al. 2018) being notable examples. Diving species of waterfowl have also been shown to have convergent morphology with other diving birds such as loons and grebes (McCracken et al. 1999; Clifton et al. 2018; Baumgart et al. 2021). Despite the preliminary evidence of widespread convergence among waterfowl, there is scant quantitative research on this phenomenon.

Geometric morphometrics is a powerful tool to study quantifiable patterns in phenotypic evolution. It has been underapplied in morphological studies of waterfowl, in which the skull and bill have received the most attention (Li and Clarke 2016; Olsen 2017). Other skeletal elements have only been examined for waterfowl as part of broader avian studies (Felice and O’Connor 2014; Marek and Felice 2014; Orkney et al. 2021) or in the recent examination of the tarsometatarsus by Mendoza and Gomez (2022). However, each of these studies have been limited in scope, either by focusing on a particular diet (Olsen 2017), single skeletal elements (Mendoza and Gomez, 2022), and/or using a limited landmark set which may have failed to detect existing ecological signals (Li and Clarke 2016). Further, these studies lack a robust and stable comparative framework for waterfowl as the available molecular phylogenetic trees have incongruent topologies due to incomplete taxon sampling, incomplete locus coverage or both (McCracken 1999; Donne-Grousse et al. 2002; Gonzalez et al. 2009; Sun et al. 2017). The need for a well-supported phylogenetic hypothesis for this research cannot be understated as the tree topology necessarily impacts detectable signals of convergence.

Integration and modularity have been an increasing focus of morphometric studies (Goswami 2007; Klingenberg and Marugan-Lobon 2014; Felice and Goswami 2018; Sherratt and Kraatz 2023). These studies have largely investigated the level of integration within a single element, often the skull (Goswami 2006; Kulemeyer et al. 2009; Piras et al 2014; Felice and Goswami et al. 2018) and have asked how fast these modules evolve in comparison to each other (Felice and Goswami 2018; Felice et al 2021; Coombs et al 2022). Few studies have investigated integration between elements, such as the skull and the appendicular skeleton (Martin-Serra et al. 2015; Randau and Goswami et al. 2018; Michaud et al. 2020; Law et al. 2024). When considering the evolution of dietary ecology in a taxonomic group, most studies have examined the skull (Foth et al. 2017; Morris et al. 2018, Felice et al. 2018; Chatterji et al. 2022; Law et al. 2022). However, studies of many groups have shown that other skeletal elements, such as the spine and limbs, show similarly strong ecological signals (Spencer 1995; Ward et al. 2002; Randau et al. 2016; Iijima and Kubo 2019; Donatelli et al. 2021). Research on the skull and appendicular elements in carnivores have shown a correlation between the degree of integration and dietary ecology suggesting selection for skeletal integration (Michaud et al. 2020; Law et al. 2024). Morphometric studies of waterfowl have focused on singular elements and have not considered whether these elements are integrated. It is notable that apparent ecological shifts in waterfowl appear to be associated with changes in locomotory behaviour when acquiring food (e.g., transitions from surface swimming to walking or diving). So, to truly capture the morphological changes associated with dietary niche evolution in waterfowl the postcranial skeleton must be studied in conjunction with the skull.

In this study, our aim is to examine ecomorphological convergence within waterfowl by conducting a thorough examination of skull and hindlimb shape relative to diet and foraging behavior. While there is strong evidence of a relationship between morphology and diet for grazing species, there have been limited quantitative investigations of other dietary niches. Specifically, we test the hypothesis that morphological convergence in waterfowl is driven by independent adaptations to similar dietary niches. First, we produce a robust, time-calibrated phylogenomic hypothesis of anseriform evolutionary relationships. We then characterize skull, femur, tibiotarsus, and tarsometatarsus shape variation using geometric morphometrics and test for signals of convergence and integration among these elements. Finally, we examine rates of phenotypic evolution and identify significant rate shifts across the phylogeny. We find support for a relationship between diet and skull shape, as well as evidence for convergent evolution across lineages that share dietary niches. We also find preliminary evidence for integration across all three measured hindlimb elements with each other and with the skull. Finally, we found that terrestrial and diving lineages had consistently higher rates of phenotypic evolution across all four elements.

## MATERIALS AND METHODS

### Sampling and DNA Extraction

We used Clements et. al. (2017) taxon list of birds to identify all known extant species of Anseriformes. We additionally sampled toe pads from two extinct waterfowl taxa, *Mergus australis* and *Anas oustaleti*. In total, we obtained 149 genetic samples from bird skin and tissue collections at multiple natural history museums (see Table S1). We isolated DNA from muscle tissue using high-throughput phenol-chloroform extraction on an AutoGenPrep 965 robot at the Smithsonian National Museum of Natural History. DNA was then dried down on the instrument, shipped, and rehydrated at Louisiana State University. We isolated DNA from toe pad specimens at the University of Oklahoma Laboratories of Molecular Anthropology and Microbiome Research (LMAMR) under strict contamination prevention control measures using a modified ancient DNA extraction following Dabney et al. (2013). Briefly we washed toe pads with 1 ml of 0.5 M EDTA for 15 minutes and then incubated at room temperature with 1 ml of 0.5 M EDTA. After 24 hours under agitation, 100 ul of Proteinase K (Qiagen) was added and the samples were incubated for several days at room temperature under agitation. After two days without progress in digestion, we incubated samples at 56°C for one hour and then continued digestion at room temperature under agitation overnight. DNA was isolated using 15 ml of guanidine hydrochloride on a Qiagen MinElute PCR Purification column. DNA was eluted twice in 30 ml of EB buffer (Qiagen).

### Library Preparation, Enrichment and Sequencing

For tissues, we prepared genomic libraries from DNA extracts for target enrichment largely following the protocol from Salter et al. (2020). First, we sheared tissue extracts using a Qsonica ultrasonicator to a fragment size distribution centered at 400-600 base pairs. We prepared dual-indexed libraries with custom indexes (Glenn et al. 2019) and the Kapa Biosystems Hyper Prep Plus library preparation kit (F. Hoffmann-La Roche AG, Basel, Switzerland) following the manufacturer’s protocol at one-half volume.

We prepared dual-indexed toe pad libraries with custom indexes (Glenn et. al. 2019) at the LMAMR following a modified protocol for degraded DNA from Caroe et al. (2017). We chose this protocol and bypassed shearing due to the DNA degradation typical of toe pad samples (McCormack et al. 2016). Briefly, we performed a partial UDG treatment on all toe pad extracts, followed by end-repair and adapter ligation. We performed qPCR on all toe pad preps to determine the optimal number of cycles per sample for the library amplification and indexing that followed using KapaHifi + Uracil following the manufacturer’s protocol (F. Hoffmann-La Roche AG, Basel, Switzerland).

We combined libraries in groups of seven or eight for a total of 18 tissue pools and three toe pad pools for enrichment. We enriched all libraries using the myBaits UCE Tetrapods 5Kv1 kit (Daciel Arbor Biosciences, Ann Arbor, MI), optimized for birds and with improved tiling for historical or ancient sources of avian DNA, following the manufacturer’s protocol. We amplified the enriched DNA pools with 16-18 cycles of PCR followed by adapter-dimer removal using a Qiagen GeneRead Size Selection kit. We then confirmed the absence of primer-dimers and verified peak size distributions on a Bioanalyzer (Agilent Technologies, Santa Clara, CA). Finally, we quantified validated, enriched pools with the KAPA qPCR quantification kit so we could produce a final equimolar pool for 150 bp paired-end sequencing at Novogene on Illumina HiSeq 2500.

### Data Quality Control and Processing

We followed the phyluce protocol for UCE phylogenomics using Phyluce 1.7.1 (Faircloth 2016). First, we removed adapter contamination and low-quality bases from raw FASTQ files using illumiprocessor (Bolger et al. 2014; Faircloth 2013; Del Fabbro et al. 2013). We then assembled the data using SPAdes (Prjibelski et al., 2020). We confirmed the taxon identification, and lack of contamination, for all sampled individuals by running raw reads corresponding to mitochondrial cytochrome oxidase I (COI) against the BOLD database (*phyluce_match_contigs_to_barcodes*) using a *Chauna torquata* COI sequence (NCBI JQ174410.1) as a reference. For species without representation in BOLD, we checked that the closest match was taxonomically coherent. To confirm that samples were not contaminated, we verified that each species set of contigs matched to only one species in BOLD. We excluded samples with inconclusive identifications or apparently contaminating sequences from the remainder of the study. We then used the Tetrapods UCE 5K probe set to bioinformatically extract UCE loci from the assembly (*phyluce_assembly_match_contigs_to_probes*). From these results, we created incomplete matrices for each locus from all processed samples, that we then aligned using mafft (Katoh et al. 2002) with default parameters. We then edge-trimmed the alignments before creating a concatenated matrix of all loci with ≥75% taxon completeness.

### Phylogenetic Analysis and Divergence Time Estimation

We ran a partitioned analysis on our concatenated dataset in IQTREE 2.0.3 (Chernomor et al. 2016; Minh et al., 2020) to estimate a maximum likelihood tree topology. We used ModelFinder (Kalyaanamoorthy et al. 2017) as implemented in IQTREE to sort individual UCE loci into an appropriate substitution model partitioning scheme. We enabled the relaxed hierarchical clustering algorithm and set the analysis to examine only the top ten percent of partition merging schemes. To assess support for the tree reconstruction, we performed 1000 replicates of ultrafast bootstrapping (Hoang et al., 2018) as well as IQ-TREE’s implementation of the SH-like approximate likelihood ratio test (Guindon et al. 2010), also with 1000 bootstrap replicates.

We also estimated the tree topology using a coalescent-based approach, SVDquartets (Chifman and Kubatko 2014; 2015), as our dataset includes historical samples (toe pads) that typically produce sequence data with fewer loci and shorter contigs relative to data generated from tissue samples (see Table S1). Prior studies suggest that SVDquartets may be less sensitive to this type of data than other tree reconstruction methods (Hosner et al. 2016; Sayyari et al. 2017). We ran our SVDquartets analysis in PAUP* v4.0a168 (Swofford 2003) with screamers (family Ahimidae) set as the outgroup, evaluating all possible quartets, and standard bootstrapping (SVDQuartets speciesTree=no bootstrap=standard evalQuartets=all nthreads=auto). We exported a majority rule consensus tree with a threshold of fifty percent.

We found discrepancies between our maximum likelihood and coalescent-based topologies. These differences were related to the placement of taxa represented by toe pad samples that were drawn together in a single clade in the maximum likelihood tree (Fig. S1). Variations of this “toepad effect” have been reported in other avian studies and have been linked to the frequent absence of the most phylogenetically informative portions of UCE flanking regions from toe pad generated data (Salter et. al. 2022). This effect was not observed in the SVDquartets tree. To probe this further, we produced maximum likelihood topologies from subsets of the concatenated alignment for clades containing toe pad samples that were placed differently between the two topologies – tribe Mergini, *Anas*, and *Hymenolaimus*. That is, we ran tree reconstruction for these clades excluding toe pads from other clades (see Fig. S2). We then combined the resulting subsampled topologies using a matrix representation with parsimony supertree approach implemented in the R-package phangorn to produce our final maximum likelihood topology (Fig. 1). We also produced a maximum likelihood topology excluding all toe pads with differential placement between analyses and used this topology for the time-tree analyses (Fig. S3), as we judged this to be the most reliable dataset for our analyses of morphological convergence (see below).

**Figure 1.**
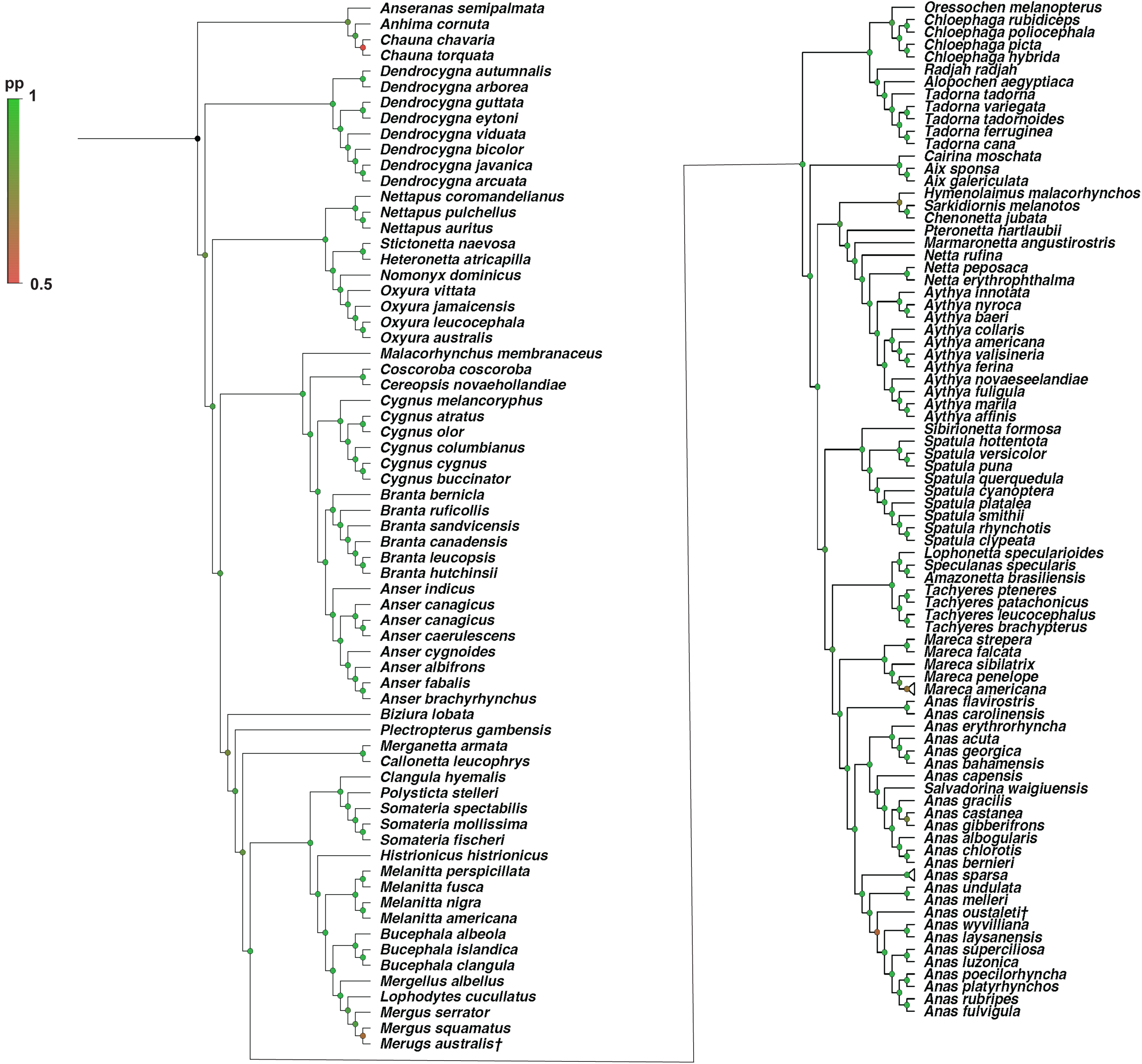
Matrix representation with parsimony supertree assembled from maximum likelihood topologies treating “wandering” toe pads separately (see Fig. S2). Circles at nodes are colored according to bootstrap support. Green indicates bootstrap values >80 as indicated in the legencd. *A dagger (†) indicates an extinct species*.

Finally, we estimated divergence times for waterfowl using BEAST 2 (Bouckaert et al. 2014) and three randomly sampled subsets of fifty UCE loci. We entered each locus as a partition into BEAUti 2.6.7 and linked loci according to the partition scheme recovered from PartitionFinder during our maximum likelihood tree reconstruction (see above) under independent GTR+GAMMA substitution models. We linked a lognormal relaxed uncorrelated rates clock model (Drummond et al. 2006) and birth-death speciation tree model across all loci. We fixed the topology based on the maximum likelihood tree with “wandering” toe pad samples removed (Fig. S3) and node-calibrated the analysis with *Vegavis iaai* (Ksepka and Clarke 2015) at Anatoidea (the split between magpie goose and the Anatidae), *Cygnopterus affinis* at the split between Anserinae and Anatinae (Mayr 2009), and *Anas soporata* at the base of Anatini (Zelenkov and Kurochkin 2012). We used a lognormal prior distribution so that 97.5 % of the distribution fell between 66 – 88ma for *Vegavis iaai* (Ksepka and Clarke 2015); 29.2 – 37.8ma for *Cygnopterus affinis* (Mayr 2009, Jones et al. 2021); 11.6 – 33.9ma for *Anas soporata,* (Zelenkov and Kurochkin 2012; Mitchell et al. 2014). We summarized posterior trees as a maximum clade credibility tree with mean node ages using TreeAnnotator v2.6.7 with the first 25% of trees discarded as burn-in (Bouckaert et al. 2014).

### Geometric Morphometrics

We analysed 158 specimens representing 118 species and 52 genera sourced from the Smithsonian National Museum of Natural History (NMNH), the Florida Museum of Natural History (FMNH), and Morphosource (Table S2). For each NMNH and FMNH specimen, we digitized the skull (n = 159), femur (n = 147), tibiotarsus (n = 145), and tarsometatarsus (n = 134) using an Einscan-SP desktop 3D scanner (Einscan 2024). We prioritized analysis of left elements; if the left element was broken or missing, we mirror imaged a mesh of the complete right element using Meshlab.

We placed landmarks for geometric morphometric analysis in Checkpoint (Stratovan Corporation). For the skull, we used a modified landmark scheme from Felice and Goswami (2018) to place 46 fixed landmarks and 21 curves consisting of 313 semilandmarks (Table S3, S4, Fig. S4 A). For the hindlimb, we used modified landmark schemes from Bjarnason and Benson (2021). We placed 17 landmarks and 242 semilandmarks in 9 curves on the femur (Table S5, S6 Fig. S4 B), 8 fixed landmarks and 117 semilandmarks in 5 curves on the tibiotarsus (Table S7, S8 Fig. S4 C), and 13 landmarks and 222 semilandmarks in 9 curves on the tarsometatarsus (Table S9, S10 Fig. S4 D). We adjusted landmarks and curves to better reflect the nuances and anatomy specific to waterfowl. The semilandmarks for each curve were then resampled and placed with even spacing across the curve using the function *subsampl.inter* in the R package SURGE (Felice 2020). The landmarks were then superimposed with a Procrustes Generalised Superimposition using the function *gpagen* in the R package Geomorph v 4.06 (Adams et al. 2024). We then deleted the bilaterally symmetrical landmarks for the skull.

### Defining Ecological Variables

We categorised the dietary ecology of each species from the existing literature (Table S11) as well as *Ducks, Geese, and Swans of the world, Revised Edition* (Johnsgard 2010). Previous categorizations of waterfowl dietary ecology have either focused solely on diet (i.e., the main nutritional resources; Olsen 2015; Olsen 2017) or foraging behaviour according to the needs of the study (Lisney et al. 2013; Pecsis et al. 2017; Mendoza and Gomez 2022; Cantlay et al. 2023). However, due to the ecological complexity of many waterfowl this can flatten the diversity in both diet and foraging seen in the group. As noted by Olsen (2017), categorising on diet alone lumped together species with distinct ecologies such as the aquatic feeding *Cygnus cygnus* and the terrestrial feeding *Branta canadensis.* Therefore, we decided to specify two discrete dietary ecology variables, foraging and diet, similar to Kalisinska (2005) and Natale and Slater (2022). Foraging describes the behaviours by which each species accesses food and is split into four categories: Surface – accessing food mostly while swimming on the surface of the water (whether that be ‘dabbling’ or ‘upending’ behaviour); Terrestrial – feeding mainly on land; Diving – primarily obtaining food while fully submerged in water; Wading – feeding while walking in shallow water or on the banks of a body of water. Diet broadly describes the main nutritional sources exploited by each species and we define five categories: Strainers – species that largely eat small particulate matter strained from the water (i.e., planktonic larvae, small seeds, and algae); Herbivores – species that feed largely on leafy plants; Piscivores – species that largely eat fish; Macroinvertivores – species that mainly eat macroinvertebrates such as molluscs and crustaceans; Mixed feeders – species that consume multiple sources of food at relatively equal rates. We then combined the foraging and diet variables to form a composite third variable, the dietary “ecotype,” to represent the broader dietary niche each species inhabits. Of the twenty possible combinations of diet and foraging, nine are observed in waterfowl (Fig. 2).

**Figure 2.**
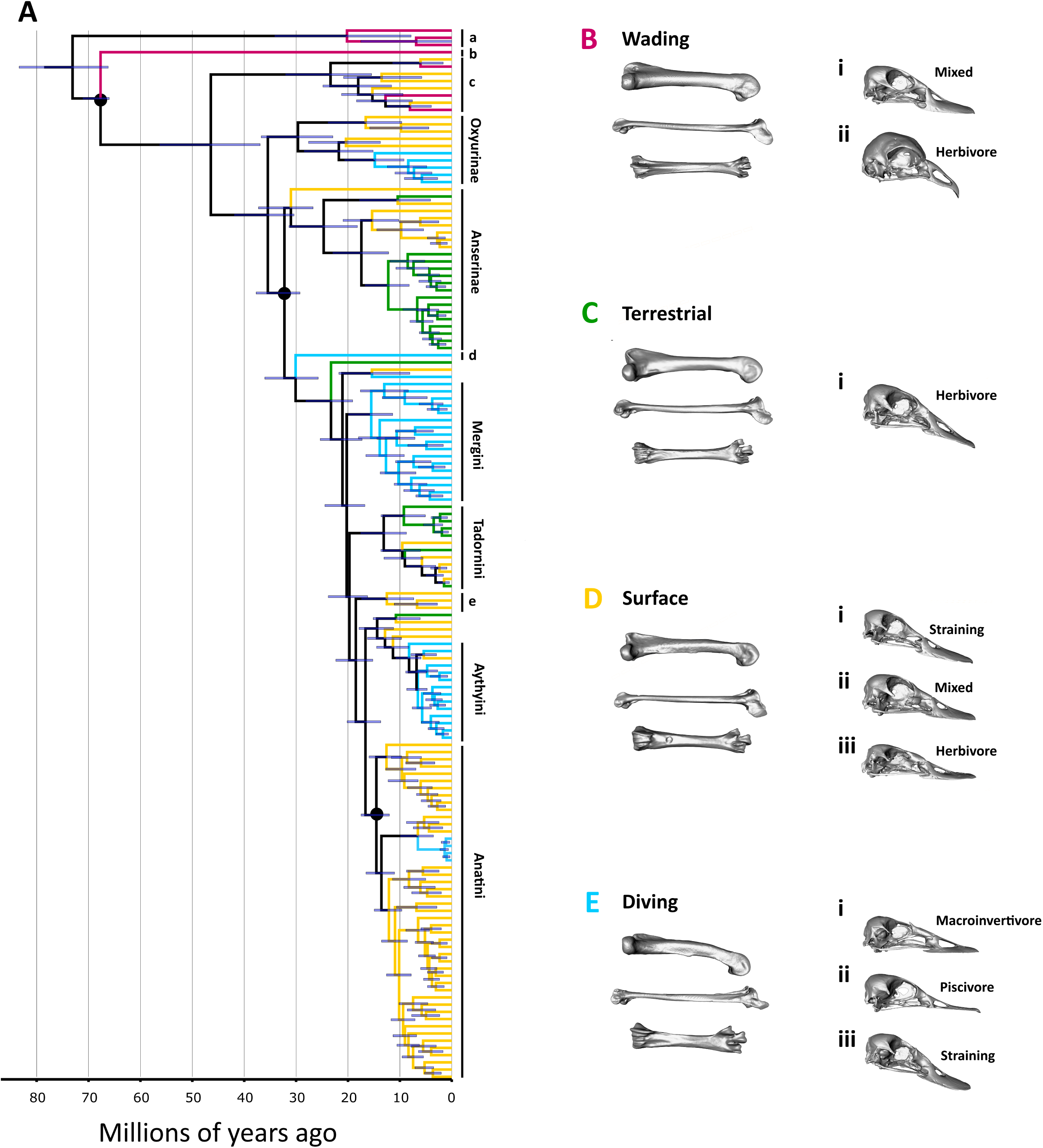
A) Time calibrated phylogeny of Anseriformes generated using Beast 2.6.7 and TreeAnnotator 2.6.7. Tips coloured based on foraging habit, purple represents wading, yellow represents surface feeding, cyan represents diving, and green represents terrestrial feeding. Node bars show 95% posterior height distribution of each node. Black circles show fossil calibration points used, from the top of figure: *Vegavis iaai, Cygnopterus affinis,* and *Anas soporata.* a is Anhimidae; b is Anseranatidae; c is Dendrocygninae; d is Biziurinae; e Cairinini. B-E; Surfaces representative of different ecological variables, with elements from species near the mean of each category. Hindlimb elements represent foraging categories and skulls represent different ecotypes. B femur is *Chauna chavaria* (NHMUK_1954.3.3), tibiotarsus is *Dendrocygna autumnalis* (NMNH 646852), tarsometatarsus is *Chauna chavaria* (NHMUK_1954.3.3). B(i) is *Dendrocygna autumnalis* (NMNH 646852). B(ii) *Chauna chavaria* (NHMUK_1954.3.3). C femur is *Branta canadensis* (NMNH 343185), tibiotarsus is *Branta canadensis* (NMNH 343185), tarsometatarsus is *Chloephaga poliocephala* (NMNH 346711). C(i) is *Anser indicus* (NMNH 557494). D femur is *Pteronetta hartlaubi* (NMNH 292390), tibiotarsus is *Anas poecilorhyncha* (NMNH 641833), tarsometatarsus is *Anas superciliosa* (NMNH 632260). D(i) is *Anas platyrhychos* (NMNH 631964). D(ii) is *Cairina moschata* (NMNH 622299). D(iii) is *Cygnus cygnus* (NMNH 430049). E femur is *Mergus serrator* (NMNH 634853), tibiotarsus is *Clangula hymenalis* (FMNH B 288708), tarsometatarsus is *Bucephala clangula* (NMNH622299). E(i) is *Somateria spectibillis* (NMNH 620823). E(ii) is *Lophodytes culculatus* (NMNH 560992). E(iii) is *Aythya ferina* (NMNH 641838).

We acknowledge that diet and ecology are complex in waterfowl and our categories serve as a reduced representation attempting to capture major features. That said, many species are behaviourally plastic and can have highly adaptable diets (Czech and Parsons 2002; Charalambidou et al. 2005; Brochet et al. 2012). We have tried to place them in categories that best capture the majority of their dietary niche, but it is important to acknowledge this simplification. Several waterfowl species have also demonstrated an ability to successfully adapt to anthropogenically modified landscapes (Czech and Parsons 2002; Brochet et al. 2012; Corriveau et al. 2020). As this adaptability has given species access to food which might not have been available historically (Czech and Parsons 2002, Hunt et al. 2019), we focused on ecological studies in largely unmodified habitats.

### Phylogenetic Comparative Methods

Phylogenetic signal was calculated using the Kmult method described in Adams (2014) using the geomorph function physignal, which is based on Blomberg’s K statistic (Blomberg et al. 2003). We pruned species without morphometric data from the time calibrated tree using the ape V5.0 (Pardis and Schliep 2019) function *drop.tip.* We then used bind.tip to add seven species that were not represented in our UCE data set but for which information on placement and divergence time was available from other prior analyses of waterfowl phylogeny (Table S12). Species without estimated divergence times were not included in further analyses.

*Testing for convergence*. – To test if one or more of the dietary variables (diet, foraging, or ecotype) had a significant impact on the morphometric evolution of the skull and hindlimb elements we employed multiple statistical analyses in R 4.3.2 (R Core Team 2024). Ornstein-Uhlenbeck models have been used widely to test for morphological convergence (Moen et al. 2016, Esquerre and Keogh 2016, Friedman et al. 2016, Grossnickle et al. 2020, Law et al. 2022). However, there have been some criticisms of this approach, including the propensity for bias towards more complex models, and sensitivity to measurement errors (Cooper et al. 2016, but see Grossnickle et al. 2023). Recent studies emphasize the need for multiple tests supporting signals of convergence, both phylogenetic comparative methods and distance-based methods (Cooper et al. 2016, Law et al. 2023, Grossnickle et al. 2023). For these reasons we employed OU models, along with C distance (Stayton 20153), and phylogenetic generalised least square (PGLS) methods.

We used the R package RPPP v1.0 to perform a phylogenetic generalised least square analysis (PGLS) with 10,000 iterations to test if the shape of each element was significantly related to size, and/or significantly related to one or more dietary variables (Collyer and Adams 2018). We fit five multivariate evolutionary models to the first five principal components (>75% of total shape variation) for each element using the R package mvMorph (Clavel et al. 2015). We ran five phylogenetic comparative models for each individual element, Brownian motion, single peak OU (OU), and three multipeak OU (OUM) models. We used a Brownian Motion model as a null model, which assumes stochastic trait variation proportional to the amount of evolutionary time and an OU model which constrains trait variation to singular optima. We then fit an OUM model for each element with each dietary variable individually (12 models total), which allowed for multiple optima defined by the assigned character state of the model variable. Ancestral states were estimated using 100 stochastically mapped trees using the function make.simmap in the R package phytools v 2.0 (Revell 2024). We assessed model fit using small sample corrected Akaike information criterion (AICc) values (Akaike 1974, Burnham and Anderson 2004). We also fit an *a priori* OUM model using the PhyloEM function of the R package PhylogeneticEM v1.6 to detect significant shifts in phenotype (Bastide et al., 2017).

Finally, we tested for morphological convergence within each dietary variable using PC distance-based methods with the R package conevol (Stayton 2015). We identified convergence using C scores, as outlined in Stayton et al. (2015).

*Testing for Integration.* – As size and corresponding allometric changes are likely strong covariables between structures in all integration analyses, we removed the component of shape correlated with size. This was done by extracting the residuals from the phylogenetic Procrustes regressions. We performed a phylogenetic two-block partial least square analysis (2B-PLS – Adams and Felice 2014) for each element pairing to test for evolutionary integration using the geomorph function phylo.integration which calculates morphological integration under a Brownian motion model of evolution.

*Evolutionary rates* – To test the effects of the dietary variables on the rate of phenotypic evolution, we used the geomorph function compare.evol.rates (Denton and Adams 2015). This function compares the net rate of change of landmark co-ordinate data along a branch under a Brownian model of evolution. The dietary variables compared for each element were those found to best support shape variation using the phylogenetic comparative models.

We also measured evolutionary rates along individual branches using the R package RRPhylo (Castiglione et al. 2018), which models heterogeneous rates of evolution and identifies shifts in phenotypic rates. The method employs phylogenetic ridge progression (Kratsh and McHardy 2014), which estimates both rates and ancestral states. The multivariate data used were PCs 1-5 for each element to keep consistent with our other phenotypic tests.

## RESULTS

### A Robust, Time-Calibrated Phylogenomic Hypothesis for Waterfowl

Table S1 summarizes the output from sequencing, assembly, and recovery of UCE loci for each taxon sample. Additional details are provided in the Supplemental Information. We produced 75% (2724 loci) ML and coalescent tree topologies (Fig. S1). Figure 1 shows the supertree topology of all sampled taxa based on the ML trees produced from the subset matrices. Results from our divergence time analyses are summarized in Figure 2.

UCE data from several galliform outgroups (not shown) confirmed the divergence between screamers (*Anhima* and *Chauna*) and the remaining anseriforms. The coalescent analysis reconstructed Oxyurinae and Anserinae as sister groups (bs=84), whereas the ML tree reconstructs these as sequentially branching (bs=100) but with the monotypic *Malacorhynchus* included as the sister taxon to Anserinae. In contrast, *Malacorhyncus* is joined with *Biziura*, another divergent monotypic lineage in our coalescent UCE topology (bs=82), and this clade is sister to the large clade of “ducks” (subfamily Anatinae). Our ML UCE topology agrees with the most comprehensively sampled mitochondrial phylogenies (Gonzalez et al., 2009; Sun et al., 2017) in placing *Biziura* on its own branch and joining *Malacorhyncus* with anserines (bs=95). Another monotypic genus, *Hymenolaimus*, similarly displays variable placement between UCE topologies, most likely due to short locus length (Table S1). In the ML tree, *Hymenolaimus* groups with *Sarkidiornis* and *Chenonetta* (bs=97; Fig. S1) in agreement with Bulgarella et al. (2010); in contrast, *Hymenolaimus* is on its own branch, diverging just after *Plectopterus* in the coalescent tree (bs=95). After the divergence of *Plectopterus*, the backbone of the coalescent topology is poorly resolved within the Anatinae, with consistently low support values (bs<70) and is best represented as a polytomy. In contrast, almost all relationships are strongly supported in the ML tree. Thus, following the consensus ML topology (Fig. 1), the order of divergence of major anseriform clades is as follows: (1) Anhimidae, (2) Anseranatidae and (3) Anatidae, representing the three recognized families within the Anseriformes and consistent with past morphological and molecular studies (e.g., Livezey 1997; Claramunt et al. 2015; Prum et al. 2015). Based on the results of our analyses and previous molecular studies, we suggest that there are six subfamily-level lineages within Anatidae, including (a) Dendrocygninae, the sister group to the rest; followed by (b) “Oxyurinae,” which includes *Nettapus* and *Stictonetta* as well as the more traditionally recognized “stifftail” ducks *Heteronetta*, *Nomonyx*, and *Oxyura*; then a clade comprising (c) Anserinae, the geese and swans, and (d) the monotypic pink-eared duck in its own subfamily “Malacorhyncinae”; followed by (e) the monotypic musk duck in its own subfamily “Biziurinae”; and finally (f) the diverse group of ducks in the Anatinae. Within Anatinae, we suggest that there are 10 lineages that merit recognition as tribes, a taxonomic rank that has been an important feature of past work on waterfowl, including (i) *Plectopterus* (ii) *Merganetta,* (iii) *Callonetta*, (iv) the sea ducks Mergini, (v) the sheldgeese and shelducks Tadornini, (vi) the perching ducks Cairinini, (vii) a clade comprising *Chenonetta*, *Sarkidiornis* and *Hymenolaimus*, (viii) a clade comprising *Pteronetta* and *Cyanochen*, (ix) the pochards and allies Aythyini, and (x) the dabbling ducks and allies Anatini. The above includes five commonly recognized tribes, albeit with modified composition in comparison to past work based on morphology (e.g., Livezey 1997), plus five additional lineages, including three that are monotypic.

The monophyly of tribes and genera are largely consistent across topologies and with previous molecular phylogenetic research (Gonzalez et al., 2009; Bulgarella et al. 2010; Fulton et al., 2012; Sun et al., 2017; Buckner et. al., 2018). Notably, *Nettapus and Stictonetta* are allied with oxyurines, and *Salvadorina* is nested within *Anas*, both results consistent with a previously unpublished genus-level analysis of waterfowl (see Fig. S5), and the latter as also suggested by morphological assessments in Delacour and Mayr (1945).

### Characterization of Skull and Hindlimb Shape Reveal Distinct Foraging Morphotypes

Figures 3 and 4 summarize the trait space for skull and hindlimb shape that characterize each ecotype and foraging mode, respectively. Skull shape is largely characterized across PC1 (35.3%) by relative bill length and skull height and across PC2 (19.4%) largely by cranio-facial angle (Fig. 3). The first PC (30.8%) of femur shape describes the degree of curvature of the femur from the midpoint of the shaft, while PC2 (24.2%) is characterised by the width of the distal femur and the relative size and position of major muscle attachments (Fig. 4a). The shape of the tibiotarsus is characterised across PC1 (41.5%) by the length of the shaft and the height of the cnemial crest; PC2 (14.02%) describes the width of the vertical blade cnemial crest (Fig. 4b). Tarsometatrsus shape is characterised across PC1 (55.7%) by its relative length and width; PC2 (9.2%) describes the relative size of the crista medialis hypotarsi and position of the trochlea (Fig. 4c). A more detailed description of the shape variation within each element is provided in the Supplemental Results.

**Figure 3.**
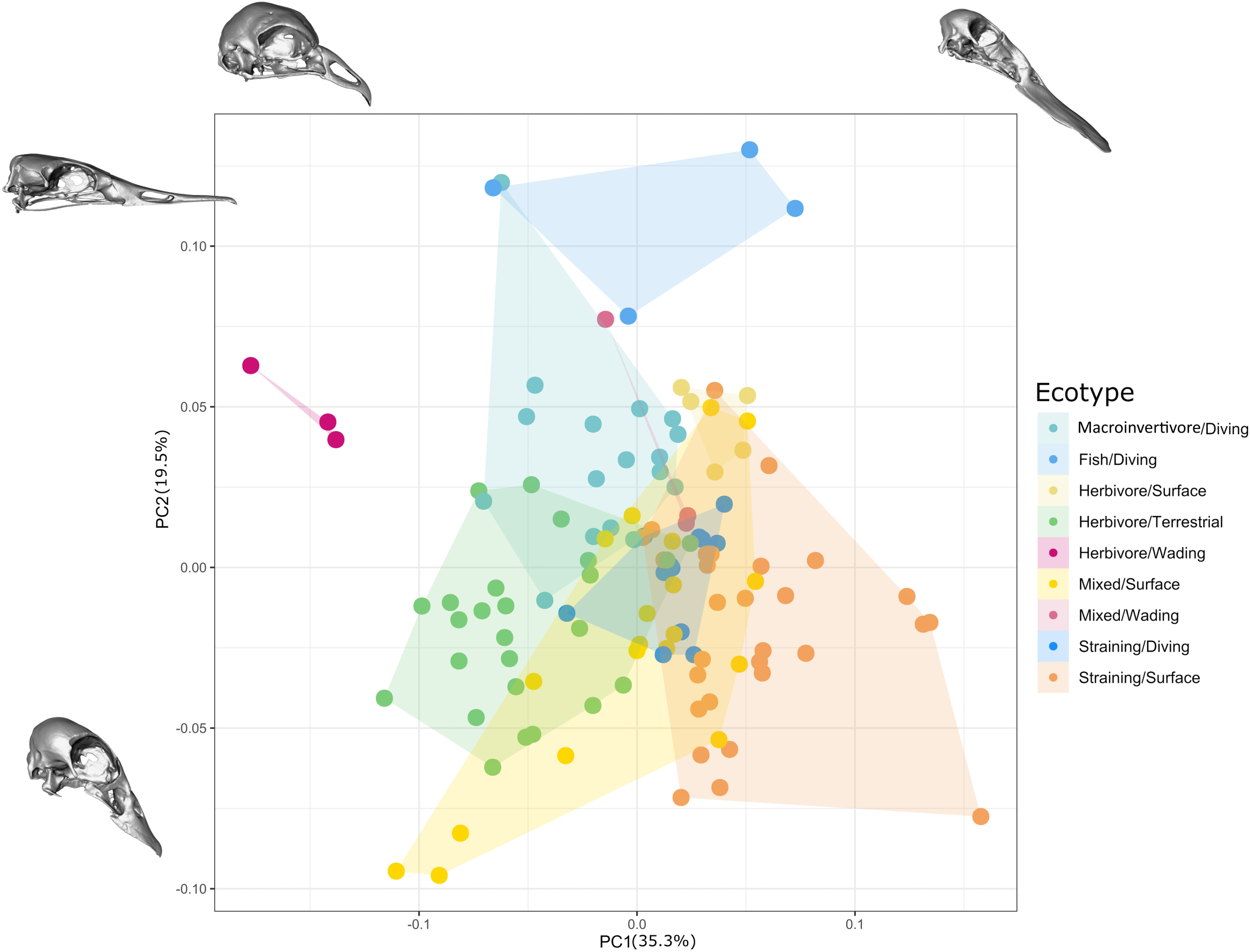
Morphospace of waterfowl skull shape showing PC1 and PC2. Different colours represent different ecotypes. Skulls shown are species with the minimum and maximum PC scores across each axis. PC1 minimum is represented by *Chauna chavaria* (NHMUK_1954.3.3). PC1 maximum is represented by *Malacororhynchus membranaceus* (BNHM 141917). PC2 minimum is represented by *Nettapus pulchellus* (NMNH 647644). PC2 maximum is represented by *Mergus squamatus* (NMNH 292752).

**Figure 4.**
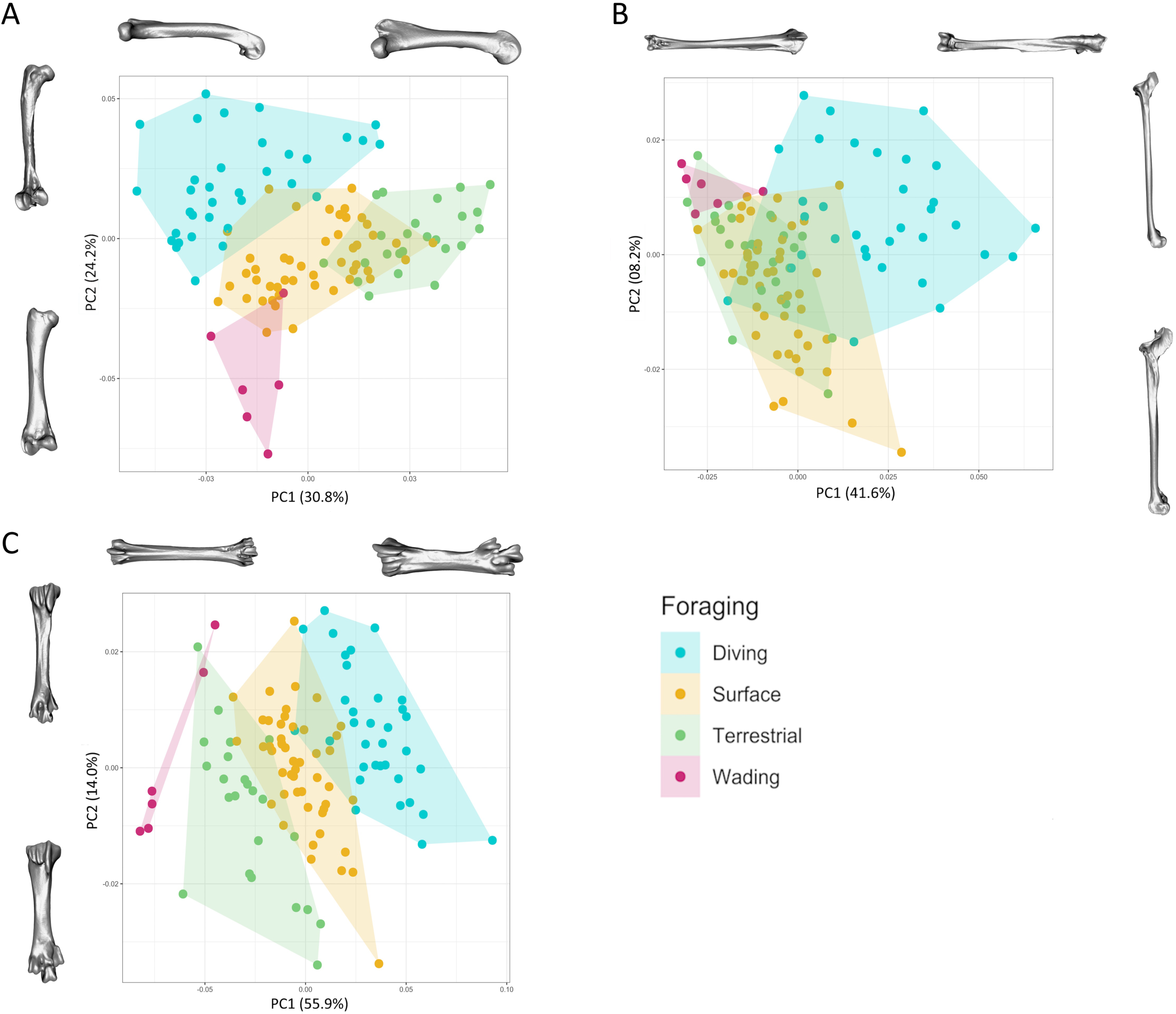
Morphospaces of waterfowl femur (A), tibiotarsus (B), and tarsometatarsus (C) shape using PC1 and PC2 for each. Surfaces shown represent species with the minimum and maximum PC scores across each axis for each element. Colours represent different foraging methods. A; PC1 minimum is represented by *Melanitta nigra* (NMNH 623466). PC1 maximum is represented by *Anser anser* (NMNH 292916). PC2 minimum is represented by *Anhima cornuta* (NMNH 226166). PC2 maximum is represented by *Melanitta fusca* (NMNH 489756). B; PC1 minimum represented by *Anhima cornuta* (NMNH 226166). PC1 maximum is represented by *Biziura lobata* (NMNH 553598). PC2 minimum is represented by *Nettapus auratus* (YPM 124151). PC2 maximum is represented by is represented by *Aythya americana* (NMNH 225501). C; PC1 minimum is represented by *Anhima cornuta* (NMNH 226166). PC1 maximum is represented by *Biziura lobata* (BNHM 143335). PC2 minimum is represented by *Anas aucklandica* (NMNH 612796). PC2 maximum is represented by *Mergus serrator* (UF-FLMNH 23708).

### Waterfowl Exhibit Strong Signals of Convergent, Diet-Driven Shape Evolution

The shape for all elements had a significant phylogenetic signal (p ≤ 0.001). The tarsometatarsus had the highest phylogenetic signal (K = 0.51) and the tibiotarsus had the lowest phylogenetic signal (K = 0.31). Results from phylogenetic least squares analyses showed that skull and tarsometatarsus shape was significantly associated with all dietary variables (P < 0.05). Femur shape was significantly related to both foraging and diet (P < 0.05), but not ecotype. Tibiotarsus shape was only significantly related to foraging habit (P = 0.03). All elements except the tibiotarsus were significantly related to size. There was a significant interaction between skull shape, size, and all dietary variables; between femur shape, size, and foraging habit (P= 0.019); and between tarsometatarsus shape, size, and diet (P = 0.047).

For all elements, the OUM was the best supported model of evolution. The ecotype OUM model best explained skull shape, whereas the foraging OUM model best explained shape variation in all hindlimb elements (Table 1; Fig. 4).

**Table 1.** Comparisons of different evolutionary models comparing different selection regimes for each of the four elements. The best fit model for each element is at the top as indicated by the smallest AICc and ΔAICc scores.

| | AICc | ΔAICc | AICcW |
| --- | --- | --- | --- |
| **Skull** |  |  |  |
| OUMecotype | -2753.8 | 0.00 | 0.99 |
| OUMdiet | -2744.5 | 9.31 | 0.01 |
| OUMforage | -2694.5 | 59.3 | 0.00 |
| BM | -2672.3 | 81.5 | 0.00 |
| OU | -2654.2 | 99.6 | 0.00 |
| **Femur** |  |  |  |
| OUMforage | -3334.0 | 0.0 | 1.00 |
| OUMecotype | -3321.0 | 13.0 | 0.00 |
| OUMdiet | -3300.5 | 33.5 | 0.00 |
| OU | -3247.8 | 86.2 | 0.00 |
| BM | -3240.7 | 93.4 | 0.00 |
| **Tibiotarsus** |  |  |  |
| OUMforage | -3731.1 | 0.0 | 1.0 |
| OUMecotype | -3697.8 | 33.3 | 0.0 |
| OUMdiet | -3675.4 | 55.7 | 0.0 |
| OU | -3673.9 | 57.2 | 0.0 |
| BM | -3658.5 | 72.6 | 0.0 |
| **Tarsometatarsus** |  |  |  |
| OUMforage | -3567.1 | 0.0 | 1.00 |
| OUMecotype | -3525.7 | 41.4 | 0.00 |
| OUMdiet | -3515.8 | 51.3 | 0.00 |
| OU | -3508.3 | 58.8 | 0.00 |
| BM | -3507.5 | 59.6 | 0.00 |

The *a priori* OUM for skull shape found 13 shifts in skull shape evolution (Fig. S6a), which largely corresponded to transitions between ecotypes. Notably, there were no detected shifts associated with diving strainers, but there were shifts within some ecotypes (e.g., adaptation to piscivory in *Mergus, Mergellus, and Lophodytes*; the characteristically extreme spatulate bills of *Spatula*). Five shifts were detected for femur shape, roughly corresponding to shifts in foraging mode (Fig. S6b). The tibiotarsus and tarsometatarsus *a priori* OUM models supported two and seven shifts, respectively (Fig. S6c-d). The detected shifts in the tibiotarsus rates occurred along the branches leading towards the Mergini and *Oxyura + Nomonyx* radiations. The shifts detected in tarsometatarsus rates occurred along the branches leading to Oxyurinae, Anserinae + Anatinae, *Bizura,* Mergini, as well as multiple individual species. Although correlations are weaker in the hindlimb than in the skull, we consistently detected shifts associated with the divergences of *Oxyura + Nomonyx*, Mergini, and *Biziura*, suggesting that diving has a strong adaptive signal in hindlimb shape.

The phenotypic distance-based methods support convergent evolution for all elements. We found support for convergence in skull shape across terrestrial herbivores (C1 = 0.164, P= 0.019), diving macroinvertivores (C1 = 0.15, P = 0.009), and surface strainers (C1 = 0.156, P < 0.001). We also found support for convergence on hindlimb element shapes across diving foragers (C1 = 0.142 – 0.242, P < 0.03) and terrestrial foragers (C1 = 0.179 – 0.248, P < 0.02). Surface strainers have convergent femur shapes (C1 = 0.124, P = 0.048), and wading species have convergent tibiotarsi (C1 = 0.197, P = 0.011) and tarsometatarsi (C1 = 0.310, P = 0.001).

***Integration.*** – We found that all elements are significantly integrated with each other (p < 0.01). The hindlimb elements are more strongly integrated with each other than any are with the skull, and all had effect sizes above 3.91 and r-PLS scores of 0.64 – 0.77. The femur and tibiotarsus have the largest effect size at 5.09 and r-PLS of 0.73. However, the skull was still significantly integrated with each hindlimb element, having the largest effect size with the tibiotarsus (Z = 3.54, r-PLS = 0.58) and a similar effect size with the femur and tarsometatarsus (Z ∼ 2.9).

***Evolutionary Rates.*** – We found that across all elements the terrestrial foraging and the dive foraging categories had the highest net rate of evolution (Fig. 5, Table S14, S15). The net evolutionary rate of the skull was significantly related to ecotype (P =0.001) and all hindlimb elements were significantly related to foraging habit (P = 0.001 for each).

**Figure 5.**
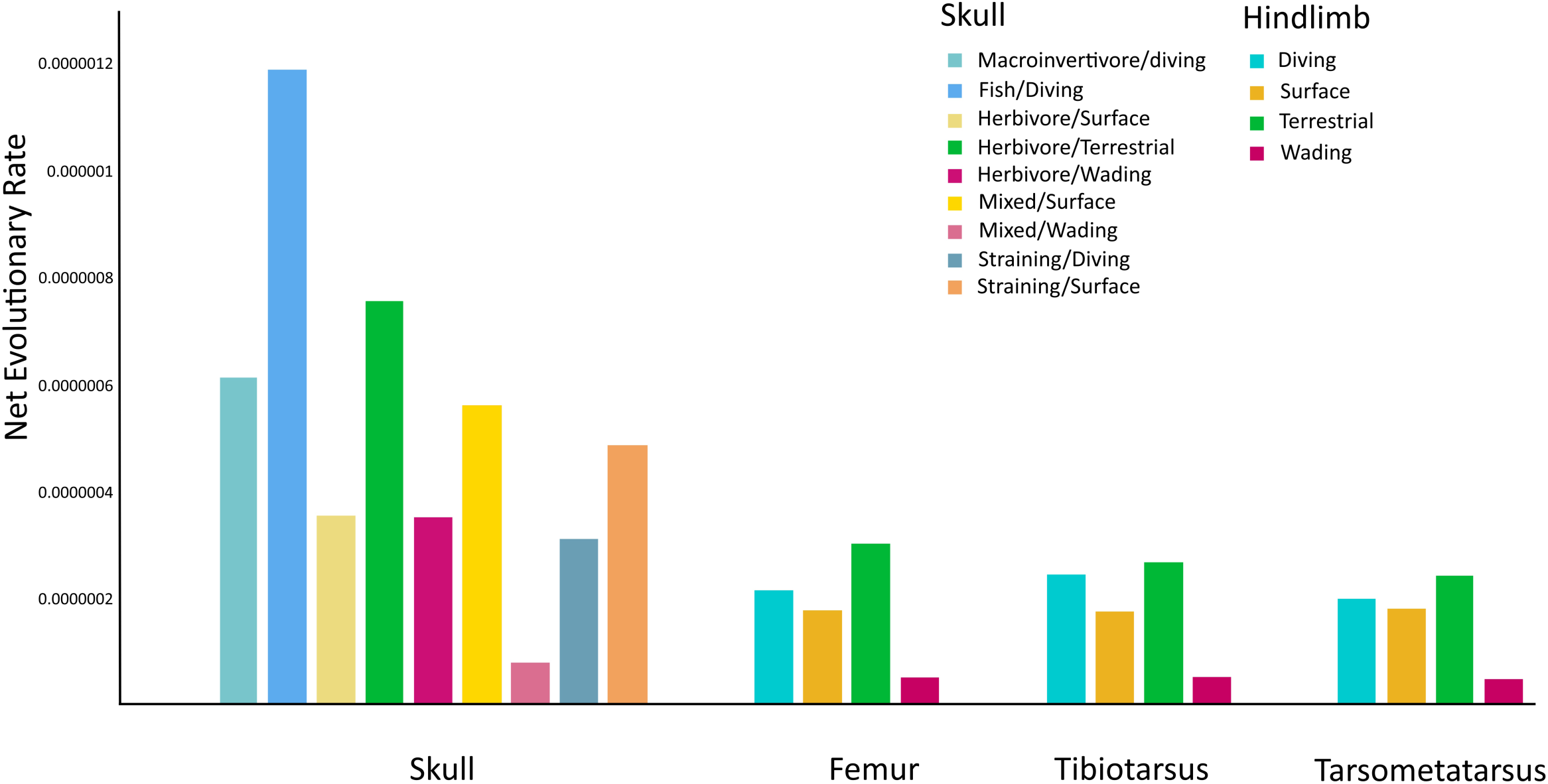
Net phenotypic evolutionary rate of each element. Rates measured for the best fitting ecological variable as based on the results of the multivariate evolutionary models for each element. Different colours represent different ecological variables.

Though we found significant shifts in the evolutionary rates of skeletal shape, they did not obviously correspond to any changes in dietary variables (Fig. S7). We detected a significant increase and a significant decrease in rates of skull shape evolution at the base of *Anas* and at the base of *Aythyini,* respectively. We found a significant rate increase in both femur and tibiotarsus shape evolution at the base of “true geese” (*Anser + Branta)*; and a significant rate increase in tarsometatarsus shape evolution at the base of Anatini excluding *Spatula + Sibirionetta formosa*).

## DISCUSSION

### A Robust, Time-Calibrated Phylogenomic Hypothesis for Waterfowl

The phylogenomic results presented are a significant step forward in waterfowl molecular systematics, representing the most comprehensively sampled and best-supported topology currently available. We did not obtain genomic data for a few notable taxa, including *Thalassornis*, *Neochen jubatus*, *Cyanochen* and *Asarcornis*, but the placement of these species has been well-resolved in previous molecular studies (Bulgarella et al. 2014; Fig. S5). Similarly, limited sequence data for *Hymenolaimus* resulted in variable placement in our analyses; however, the maximum likelihood reconstruction is more consistent with past molecular work (Bulgarella et al. 2014, Fig. S5), but future studies should further test its apparent sister relationship to *Sarkidiornis + Chenonetta*. The branch lengths toward the backbone of the tree are frequently short and likely related to the topological discordance observed between the ML and coalescent topologies. Several processes, including incomplete lineage sorting and hybridization, may explain this pattern. Modern lineages of waterfowl are known to hybridize (Lavretsky et al. 2014; Ottenburghs 2016; Lavretsky et al. 2021), which may have been an ongoing process during their early evolutionary history. Our analyses confirm that *Malacorhynchus* and *Biziura* each represent highly divergent waterfowl lineages with no close relatives and tentatively resolve their placements as sister to Anserinae and sister to Anatinae, respectively. In the future, genomic studies, perhaps coupled with filtering to capture phylogenetic signal will further clarify phylogenetic hypotheses for waterfowl, as shown for other challenging short internodes in bird phylogenetics (Gilbert et al. 2018). However, our analyses clearly demonstrate that distantly related lineages are convergent in morphology and ecology, as all topological analyses imply convergence across various clades of dabbling, diving and grazing waterfowl consistent with previous research.

We find that major clades such as Oxyurinae and Anserinae diverged during the Oligocene, earlier than some previous estimates (Gonzalez et al. 2009; Sun et al. 2017). However, this is consistent with the presence of Oligocene representatives of Oxyurinae such as *Pimpanetta* (Worthy 2009). The Early Miocene St. Bathans formation also records the presence of possible tadornines and oxyurines (Worthy et al. 2007) as well as a possible anserine (Worthy et al. 2022) suggesting that these divergences had already occurred during the Oligocene. Most major subfamilies within Anatidae (Dendrocygninae. Oxyurinae, Anserinae) are established by approximately 30Ma. This early division of major anatid clades suggests a possible undescribed fossil diversity of early anatids, and perhaps a more nested position of early genera such as *Mionetta* (Worthy and Lee 2008).

We found that the subfamily Anatinae rapidly diversified into several major groups between the Oligo-Miocene boundary (∼23Ma) and the end of the Mid-Miocene Climatic Optimum (∼14Ma) (Holbourn et al. 2015) (Fig. 4). This is largely consistent with previously dated phylogenies (Gonzalez et al. 2009; Sun et al. 2017) as well as the paleontological record (Worthy et al. 2007; Zelenkov 2020). Diversification during the Early to Mid-Miocene coincides with divergences in foraging habits, with diving and terrestrial foraging lineages arising multiple times during this period. This rapid diversification is further supported by the greater fossil anatid diversity than the earlier Oligocene record (Zelenkov and Martynovich 2013). The Early to Mid-Miocene was marked by a rise in global temperatures (Steinthorsdottir et al. 2020) which resulted in the expansion of lacustrine and wetland environments (Mandic et al. 2015; Song et al. 2018; Arenas et al. 2024). These expansions likely facilitated the establishment of modern anatid ecological diversity. By the end of the Miocene, fossil assemblages begin to record modern genera (Mlíkovský 2002, Stidham and Zelenkov 2016; Zelenkov 2020) suggesting the Miocene was a significant period for modern anatid diversification. The specific drivers of multiple ecological shifts into specialist divers and grazers, relatively rare behaviors in birds (Olsen 2015; Tyler and Younger 2022), remains unknown.

### Waterfowl Exhibit Strong Signals of Convergent, Diet-Driven Shape Evolution

We found evidence that skull and hindlimb shape is convergent across disparate groups of waterfowl and apparently corresponds to dietary niche. The considerable frequency of morphological convergence in waterfowl is strongly indicative of diet-driven adaptive evolution and contrasts evolutionary patterns reported in other avian orders (Bright et al. 2016; Sun et al. 2018; Bright et al. 2019). Foraging habit was the dominant dietary variable shaping hindlimb morphology in Anseriformes, and combined with diet formed the primary selective regime for skull shape. Ecomorphological convergence was most prominent among divers and terrestrial herbivores as suggested in previous studies (McCracken et al. 1999, Olsen 2017; Mendoza and Gomez 2022). These foraging categories also consistently displayed higher rates of phenotypic evolution across all elements. The elevated evolutionary rates likely reflect divergences from the anatid ancestral condition of surface feeding strainers (Zelenkov 2020; Musser and Clarke 2024).

Most of the convergent morphological features across the skeletal elements have clear functional significance for adaptation to a dietary niche. For example, the deeper and shorter skull of terrestrial herbivores and diving invertivores gives a mechanical advantage for applying force at the tip of the bill (Olsen 2017), whereas the shorter tarsometatarsus and proximal placement of the II trochlea are both associated with generating increased thrust in foot propelled divers (Storer 1971; McCracken et al. 1999;). While not necessarily convergent, other ecotypes show distinct adaptive morphologies, such as the long narrow bills characteristic of the mergansers which is similar to other avian piscivores (McCurry et al. 2017; Page and Cooper 2017).

Observed axes of morphological convergence across the waterfowl skull and hindlimb are integrated, consistent with evidence of widespread skeletal integration across the avian radiation (Orkney et al., 2022). Despite significant integration, the selective variables driving shape in the skull versus the hindlimb elements differ slightly. We do not think this indicates decoupled selection as proposed for carnivorans (Law et al. 2022; Law et al. 2024) because all examined shape variation is consistently correlated with foraging habit. For example, the three herbivorous ecotypes (surface, wading, and terrestrial) have drastically different skull shapes which reflect their distinct foraging behaviours and hindlimb morphologies. The long skull of surface herbivores (*Cygnus*) increases their reach when upending and searching for benthic aquatic plants (Johnsgard 2010), and the short length and tall height of the skull in terrestrial herbivores aid in grazing tougher terrestrial plants (Olsen 2017). This example illustrates the importance of jointly examining integrated characters, and multiple axes of ecological variation, to accurately capture adaptive signals (Martin-Serra et al. 2015; Michaud et al. 2020; Orkney et al. 2022; Orkney and Hendrick 2024).

We found that significant shifts in the rate of phenotypic evolution did not correspond to dietary variables, as seen in other vertebrates (labrid fish – Evans et al. 2023; muroid rodents – Alhajeri and Steppan 2018). Instead, rates of phenotypic evolution are generally high across all elements in Anatidae (Cooney et al. 2017; Felice and Goswami 2018). These high phenotypic rates mirror the numerous transitions between dietary niches, particularly during the Miocene radiation of anatines (see above). As we have shown that skull and hindlimb morphology is diet-driven, high phenotypic rates suggest ecological lability throughout waterfowl evolutionary history. Although such phenotypic lability may seem incongruent with the low morphological disparity characterizing waterfowl, we argue that their combination may contribute to the high incidence of convergence observed in across the group.

Convergence in waterfowl may largely stem from phylogenetic constraints on body morphology. Specifically, selection for the ancestral anatid condition – aquatic filter-feeders (Livezey 1997; Worthey et al. 2007; Zelenkov 2020; Musser and Clarke 2024) – drove significant morphological specializations (e.g., webbed feet, the iconic duck bill, laminae, etc.) which are largely maintained across waterfowl. Specialization can lead to reduced morphological disparity due to functional constraints on the traits (Holliday and Steppan 2004; Figueirido et al. 2016). Integration can further constrain morphology. Although the skeleton exhibits some modularity driven by distinct selection pressures, various skeletal elements must operate as a unit to achieve vital functions (e.g., procuring food and eating). This requirement to maintain both modular and integrated functions can limit the ways traits can change (Eble 2004; Felice et al. 2018), such that the same adaptive responses to selection toward niche expansion or switching is more likely to evolve than novel solutions (Sidlauskas 2008; Goswami et al. 2014). That is, waterfowl are evolutionarily confined to a restricted region of trait space (Felice et al. 2018), reducing the number of attainable adaptive optima. Thus, constrained morphological disparity, integration, and high ecological lability (evidenced by high rates of dietary niche evolution) has led to a strong pattern of morphological convergence in waterfowl.

## Conclusion

In this study, we provide the most comprehensive, best supported phylogenomic hypothesis to date for waterfowl (order Anseriformes). We then combined geometric morphometrics and recently developed phylogenetic comparative methods to demonstrate that waterfowl dietary ecology has driven convergent phenotypic evolution across multiple integrated skeletal structures. We argue that low morphological disparity, skeletal integration, and ecological lability help explain the propensity for phenotypic convergence in this group. Our results indicate that waterfowl are a promising model group for continued studies of the causes and consequences of adaptive evolutionary processes.

## Acknowledgements

The authors would like to acknowledge the following sources of funding: NSF DBI-1711968 to JCB; and NSF-DEB 0089760 to MDS and HFJ. The authors thank Brant Faircloth for assistance with data collection and analysis via the “OpenWings” project (NSF DEB-1655559, DEB-1655624, DEB-1655683, DEB-1655736). We also thank Chris Abin, Nihan Dagtas, Karissa Hughes, and Glaucia Del-Rio for assistance with laboratory work. We acknowledge Morphosource and its contributors for data access. We especially acknowledge and thank the curators and collection managers, including Andrew Kratter, Christopher Milensky, Libby Beckman, Joel Cracraft, Paul Sweet, Thomas Trombone, Stephen Rogers, David Winkler, Irby Lovette, Ben Marks, Shannon Hacket, John Bates, Frederick Sheldon, Robb Brumfield, Donna Dittmann, Keith Barker, Christopher Witt, Kevin Winker, Christopher Milensky, Sharon Birks, and Kristof Zyskowski, who generously granted access to specimens, sub-sampled and shipped the tissues that made this research possible.

## References

1. Adams D, Collyer M, Kaliontzopoulou A, Baken E (2024). “Geomorph: Software for geometric morphometric analyses. R package version 4.0.8.” https://cran.r-project.org/package=geomorph.

2. Adams, D. C. (2014). A Generalized K Statistic for Estimating Phylogenetic Signal from Shape and Other High-Dimensional Multivariate Data. Systematic Biology, 63(5), 685– 697. 10.1093/sysbio/syu030

3. Adams, D. C., & Felice, R. N. (2014). Assessing Trait Covariation and Morphological Integration on Phylogenies Using Evolutionary Covariance Matrices. PLOS ONE, 9(4), e94335. 10.1371/journal.pone.0094335

4. Akaike, H. (1974). A new look at the statistical model identification. IEEE Transactions on Automatic Control, 19(6), 716–723. IEEE Transactions on Automatic Control. 10.1109/TAC.1974.1100705

5. Alhajeri, B. H., & Steppan, S. J. (2018). Disparity and Evolutionary Rate Do Not Explain Diversity Patterns in Muroid Rodents (Rodentia: Muroidea). Evolutionary Biology, 45(3), 324–344. 10.1007/s11692-018-9453-z

6. Arenas, C., Osácar, C., Pérez-Rivarés, F. J., Bastida, J., Gil, A., & Auqué, L. F. (n.d.). The Early–Middle Miocene climate as reflected by a mid-latitude lacustrine record in the Ebro Basin, north-east Iberia. The Depositional Record, n/a(n/a). 10.1002/dep2.290

7. Baeckens, S., Goeyers, C., & Van Damme, R. (2020). Convergent Evolution of Claw Shape in a Transcontinental Lizard Radiation. Integrative and Comparative Biology, 60(1), 10–23. 10.1093/icb/icz151

8. Bastide, P., Mariadassou, M., & Robin, S. (2017). Detection of Adaptive Shifts on Phylogenies by using Shifted Stochastic Processes on a Tree. Journal of the Royal Statistical Society Series B: Statistical Methodology, 79(4), 1067–1093. 10.1111/rssb.12206

9. Baumgart, S. L., Sereno, P. C., & Westneat, M. W. (2021). Wing Shape in Waterbirds: Morphometric Patterns Associated with Behavior, Habitat, Migration, and Phylogenetic Convergence. Integrative Organismal Biology, 3(1), obab011. 10.1093/iob/obab011

10. Blomberg, S. P., Garland JR., T., & Ives, A. R. (2003). Testing for Phylogenetic Signal in Comparative Data: Behavioral Traits Are More Labile. Evolution, 57(4), 717–745. 10.1111/j.0014-3820.2003.tb00285.x

11. Bjarnason, A., & Benson, R. B. J. (2021). A 3D geometric morphometric dataset quantifying skeletal variation in birds. MorphoMuseuM, 7(1). https://ora.ox.ac.uk/objects/uuid:96ad8bec-2c16-4278-a384-657f309c37fd

12. Bolger, A. M., Lohse, M., & Usadel, B. (2014). Trimmomatic: A flexible trimmer for Illumina Sequence Data. Bioinformatics. 10.1093/bioinformatics/btu170.

13. Bouckaert R, Heled J, Kühnert D, Vaughan T, Wu CH, Xie D, et al. BEAST 2: a software platform for Bayesian evolutionary analysis. PLoS computational biology. 2014;10(4):e1003537. pmid:24722319

14. Bouckaert R, Vaughan TG, Barido-Sottani J, Duchêne S, Fourment M, Gavryushkina A, et al. (2019) BEAST 2.5: An advanced software platform for Bayesian evolutionary analysis. PLoS Comput Biol 15(4): e1006650. 10.1371/journal.pcbi.1006650

15. Brochet, A.-L., Dessborn, L., Legagneux, P., Elmberg, J., Gauthier-Clerc, M., Fritz, H., & Guillemain, M. (2012). Is diet segregation between dabbling ducks due to food partitioning? A review of seasonal patterns in the Western Palearctic. Journal of Zoology, 286(3), 171–178. 10.1111/j.1469-7998.2011.00870.x

16. Buckner, J. C., Ellingson, R., Gold, D. A., Jones, T. L., & Jacobs, D. K. (2018). Mitogenomics supports an unexpected taxonomic relationship for the extinct diving duck Chendytes lawi and definitively places the extinct Labrador Duck. Molecular Phylogenetics and Evolution, 122, 102–109. 10.1016/j.ympev.2017.12.008

17. Bulgarella, M., Kopuchian, C., Giacomo, A. S. D., Matus, R., Blank, O., Wilson, R. E., & Mccracken, K. G. (2014). Molecular phylogeny of the South American sheldgeese with implications for conservation of Falkland Islands (Malvinas) and continental populations of the Ruddy-headed Goose Chloephaga rubidiceps and Upland Goose C. picta. Bird Conservation International, 24(1), 59–71. 10.1017/S0959270913000178

18. Burnham, K. P., & Anderson, D. R. (2004). Multimodel Inference: Understanding AIC and BIC in Model Selection. Sociological Methods & Research, 33(2), 261–304. 10.1177/0049124104268644

19. Carøe, C., Gopalakrishnan, S., Vinner, L., Mak, S. S. T., Sinding, M. H. S., Samaniego, J. A., Wales, N., Sicheritz-Pontén, T., & Gilbert, M. T. P. (2018). Single-tube library preparation for degraded DNA. Methods in Ecology and Evolution, 9(2), 410–419. 10.1111/2041-210X.12871

20. Castiglione, S., Tesone, G., Piccolo, M., Melchionna, M., Mondanaro, A., Serio, C., Di Febbraro, M., & Raia, P. (2018). A new method for testing evolutionary rate variation and shifts in phenotypic evolution. Methods in Ecology and Evolution, 9(4), 974–983. 10.1111/2041-210X.12954

21. Charalambidou, I., Santamaría, L., Jansen, C., & Nolet, B. A. (2005). Digestive Plasticity in Mallard Ducks Modulates Dispersal Probabilities of Aquatic Plants and Crustaceans. Functional Ecology, 19(3), 513–519.

22. Chatterji, R. M., Hipsley, C. A., Sherratt, E., Hutchinson, M. N., & Jones, M. E. H. (2022). Ontogenetic allometry underlies trophic diversity in sea turtles (Chelonioidea). Evolutionary Ecology, 36(4), 511–540. 10.1007/s10682-022-10162-z

23. Chernomor, O., von Haeseler, A., Minh, B.Q. (2016) Terrace aware data structure for phylogenomic inference from supermatrices. Syst. Biol., 65:997–1008. 10.1093/sysbio/syw037

24. Chifman, J., & Kubatko, L. (2014). Quartet Inference from SNP Data Under the Coalescent Model. Bioinformatics, 30(23), 3317–3324. 10.1093/bioinformatics/btu530

25. Chifman, J., & Kubatko, L. (2015). Identifiability of the unrooted species tree topology under the coalescent model with time-reversible substitution processes, site-specific rate variation, and invariable sites. Journal of Theoretical Biology, 374, 35–47. 10.1016/j.jtbi.2015.03.006

26. Clements, J. F., T. S. Schulenberg, M. J. Iliff, D. Roberson, T. A. Fredericks, B. L. Sullivan, and C. L. Wood. 2017. The eBird/Clements checklist of birds of the world: v2016. Downloaded from https://www.birds.cornell.edu/clementschecklist/august-2017/2017-citation-checklist-download/

27. Clavel, J., Escarguel, G., & Merceron, G. (2015). mvmorph: An r package for fitting multivariate evolutionary models to morphometric data. Methods in Ecology and Evolution, 6(11), 1311–1319. 10.1111/2041-210X.12420

28. Clifton, G. T., Carr, J. A., & Biewener, A. A. (2018). Comparative hindlimb myology of foot-propelled swimming birds. Journal of Anatomy, 232(1), 105–123. 10.1111/joa.12710

29. Coombs, E. J., Felice, R. N., Clavel, J., Park, T., Bennion, R. F., Churchill, M., Geisler, J. H., Beatty, B., & Goswami, A. (2022). The tempo of cetacean cranial evolution. Current Biology, 32(10), 2233–2247.e4. 10.1016/j.cub.2022.04.060

30. Cooney, C. R., Bright, J. A., Capp, E. J. R., Chira, A. M., Hughes, E. C., Moody, C. J. A., Nouri, L. O., Varley, Z. K., & Thomas, G. H. (2017). Mega-evolutionary dynamics of the adaptive radiation of birds. Nature, 542(7641), 344–347. 10.1038/nature21074

31. Cooper, N., Thomas, G. H., Venditti, C., Meade, A., & Freckleton, R. P. (2016). A cautionary note on the use of Ornstein Uhlenbeck models in macroevolutionary studies. Biological Journal of the Linnean Society, 118(1), 64–77. 10.1111/bij.12701

32. Corriveau, A., Klaassen, M., Crewe, T. L., Kaestli, M., Garnett, S. T., Loewensteiner, D. A., Rogers, R. M., & Campbell, H. A. (2020). Broad-scale opportunistic movements in the tropical waterbird Anseranas semipalmata: Implications for human-wildlife conflicts. Emu – Austral Ornithology, 120(4), 343–354. 10.1080/01584197.2020.1857651

33. Czech, H. A., & Parsons, K. C. (2002). Agricultural Wetlands and Waterbirds: A Review. Waterbirds: The International Journal of Waterbird Biology, 25, 56–65.

34. Dabney, J., Knapp, M., Glocke, I., Gansauge, M.-T., Weihmann, A., Nickel, B., Valdiosera, C., García, N., Pääbo, S., Arsuaga, J.-L., & Meyer, M. (2013). Complete mitochondrial genome sequence of a Middle Pleistocene cave bear reconstructed from ultrashort DNA fragments. Proceedings of the National Academy of Sciences, 110(39), 15758–15763. 10.1073/pnas.1314445110

35. De Mendoza, R. S., & Gómez, R. O. (2022). Ecomorphology of the tarsometatarsus of waterfowl (Anseriformes) based on geometric morphometrics and its application to fossils. The Anatomical Record, 305(11), 3243–3253. 10.1002/ar.24891

36. Del Fabbro, C. D., Scalabrin, S., Morgante, M., & Giorgi, F. M. (2013). An Extensive Evaluation of Read Trimming Effects on Illumina NGS Data Analysis. PLOS ONE, 8(12), e85024. 10.1371/journal.pone.0085024

37. Denton, J. S. S., & Adams, D. C. (2015). A new phylogenetic test for comparing multiple high-dimensional evolutionary rates suggests interplay of evolutionary rates and modularity in lanternfishes (Myctophiformes; Myctophidae). Evolution, 69(9), 2425– 2440. 10.1111/evo.12743

38. Donatelli, C. M., Roberts, A. S., Scott, E., DeSmith, K., Summers, D., Abu-Bader, L., Baxter, D., Standen, E. M., Porter, M. E., Summers, A. P., & Tytell, E. D. (2021). Foretelling the Flex—Vertebral Shape Predicts Behavior and Ecology of Fishes. Integrative and Comparative Biology, 61(2), 414–426. 10.1093/icb/icab110

39. Donne-Goussé, C., Laudet, V., & Hänni, C. (2002). A molecular phylogeny of anseriformes based on mitochondrial DNA analysis. Molecular Phylogenetics and Evolution, 23(3), 339–356. 10.1016/S1055-7903(02)00019-2

40. Drummond, A. J., Ho, S. Y. W., Phillips, M. J., & Rambaut, A. (2006). Relaxed Phylogenetics and Dating with Confidence. PLOS Biology, 4(5), e88. 10.1371/journal.pbio.0040088

41. Eble, G. J. (2004). The macroevolution of phenotypic integration. Phenotypic Integration: Studying the Ecology and Evolution of Complex Phenotypes, 253–273.

42 EinScan Multifunctional 3D Scanner Specs. (2024, February 28). EinScan. https://www.einscan.com/einscan-sp/einscan-sp-specs/

43. Esquerré, D., & Scott Keogh, J. (2016). Parallel selective pressures drive convergent diversification of phenotypes in pythons and boas. Ecology Letters, 19(7), 800–809. 10.1111/ele.12620

44. Evans, K. M., Larouche, O., Gartner, S. M., Faucher, R. E., Dee, S. G., & Westneat, M. W. (2023). Beaks promote rapid morphological diversification along distinct evolutionary trajectories in labrid fishes (Eupercaria: Labridae). Evolution, 77(9), 2000–2014. 10.1093/evolut/qpad115

45. Evers, S. W., & Benson, R. B. J. (2019). A new phylogenetic hypothesis of turtles with implications for the timing and number of evolutionary transitions to marine lifestyles in the group. Palaeontology, 62(1), 93–134. 10.1111/pala.12384

46. Faircloth, B. (2013). Illumiprocessor: A trimmomatic wrapper for parallel adapter and quality trimming.

47. Faircloth, B. C. (2016). PHYLUCE is a software package for the analysis of conserved genomic loci. *Bioinformatics (Oxford*, England), 32(5), 786–788. 10.1093/bioinformatics/btv646

48. Faith, D. P. (1989). Homoplasy as Pattern: Multivariate Analysis of Morphological Convergence in Anseriformes. Cladistics, 5(3), 235–258. 10.1111/j.1096-0031.1989.tb00488.x

49. Felice, R.N, SURGE, (2020), GitHub repository, https://github.com/rnfelice/SURGE

50. Felice, R. N., & Goswami, A. (2018). Developmental origins of mosaic evolution in the avian cranium. Proceedings of the National Academy of Sciences, 115(3), 555–560. 10.1073/pnas.1716437115

51. Felice, R. N., & O’Connor, P. M. (2014). Ecology and Caudal Skeletal Morphology in Birds: The Convergent Evolution of Pygostyle Shape in Underwater Foraging Taxa. PLOS ONE, 9(2), e89737. 10.1371/journal.pone.0089737

52. Felice, R. N., Tobias, J. A., Pigot, A. L., & Goswami, A. (2019). Dietary niche and the evolution of cranial morphology in birds. Proceedings of the Royal Society B: Biological Sciences, 286(1897), 20182677. 10.1098/rspb.2018.2677

53. Felice, R. N., Watanabe, A., Cuff, A. R., Noirault, E., Pol, D., Witmer, L. M., Norell, M. A., O’Connor, P. M., & Goswami, A. (2019). Evolutionary Integration and Modularity in the Archosaur Cranium. Integrative and Comparative Biology, 59(2), 371–382. 10.1093/icb/icz052

54. Figueirido, B., MacLeod, N., Krieger, J., Renzi, M. D., Pérez-Claros, J. A., & Palmqvist, P. (2011). Constraint and adaptation in the evolution of carnivoran skull shape. Paleobiology, 37(3), 490–518. 10.1666/09062.1

55. Foth, C., Rabi, M., & Joyce, W. G. (2017). Skull shape variation in extant and extinct Testudinata and its relation to habitat and feeding ecology. Acta Zoologica, 98(3), 310–325. 10.1111/azo.12181

56. Friedman, S. T., Price, S. A., Hoey, A. S., & Wainwright, P. C. (2016). Ecomorphological convergence in planktivorous surgeonfishes. Journal of Evolutionary Biology, 29(5), 965– 978. 10.1111/jeb.12837

57. Gibbs, H. L., & Grant, P. R. (1987). Ecological Consequences of an Exceptionally Strong El Nino Event on Darwin’s Finches. Ecology, 68(6), 1735–1746. 10.2307/1939865

58. Gilbert, P. S., Wu, J., Simon, M. W., Sinsheimer, J. S., & Alfaro, M. E. (2018). Filtering nucleotide sites by phylogenetic signal to noise ratio increases confidence in the Neoaves phylogeny generated from ultraconserved elements. Molecular Phylogenetics and Evolution, 126, 116–128. 10.1016/j.ympev.2018.03.033

59. Glenn, T. C., Nilsen, R. A., Kieran, T. J., Sanders, J. G., Bayona-Vásquez, N. J., Finger, J. W., Pierson, T. W., Bentley, K. E., Hoffberg, S. L., Louha, S., Leon, F. J. G.-D., Portilla, M. A. del R., Reed, K. D., Anderson, J. L., Meece, J. K., Aggrey, S. E., Rekaya, R., Alabady, M., Belanger, M., … Faircloth, B. C. (2019). Adapterama I: Universal stubs and primers for 384 unique dual-indexed or 147,456 combinatorially-indexed Illumina libraries (iTru & iNext). PeerJ, 7, e7755. 10.7717/peerj.7755

60. Gonzalez, J., Düttmann, H., & Wink, M. (2009). Phylogenetic relationships based on two mitochondrial genes and hybridization patterns in Anatidae. Journal of Zoology, 279(3), 310–318. 10.1111/j.1469-7998.2009.00622.x

61. Goswami, A. (2007). Phylogeny, Diet, and Cranial Integration in Australodelphian Marsupials. PLOS ONE, 2(10), e995. 10.1371/journal.pone.0000995

62. Goswami, A., Smaers, J. B., Soligo, C., & Polly, P. D. (2014). The macroevolutionary consequences of phenotypic integration: From development to deep time. Philosophical Transactions of the Royal Society B: Biological Sciences, 369(1649), 20130254. 10.1098/rstb.2013.0254

63. Grossnickle, D. M., Brightly, W. H., Weaver, L. N., Stanchak, K. E., Roston, R. A., Pevsner, S. K., Stayton, C. T., Polly, P. D., & Law, C. J. (2024). Challenges and advances in measuring phenotypic convergence. *Evolution*, qpae081. 10.1093/evolut/qpae081

64. Grossnickle, D. M., Chen, M., Wauer, J. G. A., Pevsner, S. K., Weaver, L. N., Meng, Q.-J., Liu, D., Zhang, Y.-G., & Luo, Z.-X. (2020). Incomplete convergence of gliding mammal skeletons*. Evolution, 74(12), 2662–2680. 10.1111/evo.14094

65. Guindon, S., Dufayard, J.-F., Lefort, V., Anisimova, M., Hordijk, W., & Gascuel, O. (2010). New Algorithms and Methods to Estimate Maximum-Likelihood Phylogenies: Assessing the Performance of PhyML 3.0. Systematic Biology, 59(3), 307–321. 10.1093/sysbio/syq010

66. Hansen, T. F., Pienaar, J., & Orzack, S. H. (2008). A comparative method for studying adaptation to a randomly evolving environment. Evolution, 62(8), 1965–1977. 10.1111/j.1558-5646.2008.00412.x

67. Hoang, D. T., Chernomor, O., von Haeseler, A., Minh, B. Q., & Vinh, L. S. (2018). UFBoot2: Improving the Ultrafast Bootstrap Approximation. Molecular Biology and Evolution, 35(2), 518–522. 10.1093/molbev/msx281

68. Holbourn, A., Kuhnt, W., Kochhann, K. G. D., Andersen, N., & Sebastian Meier, K. J. (2015). Global perturbation of the carbon cycle at the onset of the Miocene Climatic Optimum. Geology, 43(2), 123–126. 10.1130/G36317.1

69. Holliday, J. A., & Steppan, S. J. (2004). Evolution of hypercarnivory: The effect of specialization on morphological and taxonomic diversity. Paleobiology, 30(1), 108–128. 10.1666/0094-8373(2004)030<0108:EOHTEO>2.0.CO;2

70. Hosner, P. A., Faircloth, B. C., Glenn, T. C., Braun, E. L., & Kimball, R. T. (2016). Avoiding Missing Data Biases in Phylogenomic Inference: An Empirical Study in the Landfowl (Aves: Galliformes). Molecular Biology and Evolution, 33(4), 1110–1125. 10.1093/molbev/msv347

71. Iijima, M., & Kubo, T. (2019). Comparative morphology of presacral vertebrae in extant crocodylians: Taxonomic, functional and ecological implications. Zoological Journal of the Linnean Society, 186(4), 1006–1025. 10.1093/zoolinnean/zly096

72. Jaekel, M., & Wake, D. B. (2007). Developmental processes underlying the evolution of a derived foot morphology in salamanders. Proceedings of the National Academy of Sciences, 104(51), 20437–20442. 10.1073/pnas.0710216105

73. James, H. F., & Burney, D. A. (1997). The diet and ecology of Hawaii’s extinct flightless waterfowl: Evidence from coprolites. Biological Journal of the Linnean Society, 62(2), 279–297. 10.1111/j.1095-8312.1997.tb01627.x

74. Johnsgard, P. (2010). Waterfowl of North America: The Biology of Waterfowl. Waterfowl of North America, Revised Edition (2010) by Paul A. Johnsgard. https://digitalcommons.unl.edu/biosciwaterfowlna/4

75. Jones, T. L., Coltrain, J. B., Jacobs, D. K., Porcasi, J., Brewer, S. C., Buckner, J. C., Perrine, J. D., & Codding, B. F. (2021). Causes and consequences of the late Holocene extinction of the marine flightless duck (*Chendytes lawi*) in the northeastern Pacific. Quaternary Science Reviews, 260, 106914. 10.1016/j.quascirev.2021.106914

76. Katoh, K., Misawa, K., Kuma, K., & Miyata, T. (2002). MAFFT: A novel method for rapid multiple sequence alignment based on fast Fourier transform. Nucleic Acids Research, 30(14), 3059–3066. 10.1093/nar/gkf436

77. Kalisińska, E. (2005). Anseriform Brain and Its Parts versus Taxonomic and Ecological Categories. Brain Behavior and Evolution, 65(4), 244–261. 10.1159/000084315

78. Kalyaanamoorthy, S., Minh, B.Q., Wong, T.K.F., von Haeseler, A., Jermiin, L.S. (2017) ModelFinder: Fast model selection for accurate phylogenetic estimates. Nat. Methods, 14:587–589. 10.1038/nmeth.4285

79. Kess, T., & Boulding, E. G. (2019). Genome-wide association analyses reveal polygenic genomic architecture underlying divergent shell morphology in Spanish Littorina saxatilis ecotypes. Ecology and EvolAution, 9(17), 9427–9441. 10.1002/ece3.5378

80. Klingenberg, C., & Marugán-Lobón, J. (2013). Evolutionary Covariation in Geometric Morphometric Data: Analyzing Integration, Modularity, and Allometry in a Phylogenetic Context. Systematic Biology, 62. 10.1093/sysbio/syt025

81. Kratsch, C., & McHardy, A. C. (2014). RidgeRace: Ridge regression for continuous ancestral character estimation on phylogenetic trees. Bioinformatics, 30(17), i527–i533. 10.1093/bioinformatics/btu477

82. Ksepka, D., & Clarke, J. (2015). Phylogenetically vetted and stratigraphically constrained fossil calibrations within Aves. Palaeontologia Electronica, 18(1), 1–25.

83. Kulemeyer, C., Asbahr, K., Gunz, P., Frahnert, S., & Bairlein, F. (2009). Functional morphology and integration of corvid skulls – a 3D geometric morphometric approach. Frontiers in Zoology, 6(1), 2. 10.1186/1742-9994-6-2

84. Lavretsky, P., Hernández-Baños, B. E., & Peters, J. L. (2014). Rapid radiation and hybridization contribute to weak differentiation and hinder phylogenetic inferences in the New World Mallard complex (Anas spp.). The Auk, 131(4), 524–538. 10.1642/AUK-13-164.1

85. Lavretsky, P., Wilson, R. E., Talbot, S. L., & Sonsthagen, S. A. (2021). Phylogenomics reveals ancient and contemporary gene flow contributing to the evolutionary history of sea ducks (Tribe Mergini). Molecular Phylogenetics and Evolution, 161, 107164. 10.1016/j.ympev.2021.107164

86. Law, C. J., Blackwell, E. A., Curtis, A. A., Dickinson, E., Hartstone-Rose, A., & Santana, S. E. (2022). Decoupled evolution of the cranium and mandible in carnivoran mammals. Evolution, 76(12), 2959–2974. 10.1111/evo.14578

87. Law, C. J., Hlusko, L. J., & Tseng, Z. J. (2024). Uncovering the mosaic evolution of the carnivoran skeletal system. Biology Letters, 20(1), 20230526. 10.1098/rsbl.2023.0526

88. Lerner, H. R. L., Meyer, M., James, H. F., Hofreiter, M., & Fleischer, R. C. (2011). Multilocus Resolution of Phylogeny and Timescale in the Extant Adaptive Radiation of Hawaiian Honeycreepers. Current Biology, 21(21), 1838–1844. 10.1016/j.cub.2011.09.039

89. Li, Z., & Clarke, J. A. (2016). The Craniolingual Morphology of Waterfowl (Aves, Anseriformes) and Its Relationship with Feeding Mode Revealed Through Contrast-Enhanced X-Ray Computed Tomography and 2D Morphometrics. Evolutionary Biology, 43(1), 12–25. 10.1007/s11692-015-9345-4

90. Lisney, T. J., Stecyk, K., Kolominsky, J., Schmidt, B. K., Corfield, J. R., Iwaniuk, A. N., & Wylie, D. R. (2013). Ecomorphology of eye shape and retinal topography in waterfowl (Aves: Anseriformes: Anatidae) with different foraging modes. Journal of Comparative Physiology A, 199(5), 385–402. 10.1007/s00359-013-0802-1

91. Livezey, B. C. (1997). A phylogenetic analysis of basal Anseriformes, the fossil Presbyornis, and the interordinal relationships of waterfowl. Zoological Journal of the Linnean Society, 121(4), 361–428. 10.1111/j.1096-3642.1997.tb01285.x

92. Losos, J. B. (2011). CONVERGENCE, ADAPTATION, AND CONSTRAINT. Evolution, 65(7), 1827–1840. 10.1111/j.1558-5646.2011.01289.x

93. Mandic, O., de Leeuw, A., Vuković, B., Krijgsman, W., Harzhauser, M., & Kuiper, K. F. (2011). Palaeoenvironmental evolution of Lake Gacko (Southern Bosnia and Herzegovina): Impact of the Middle Miocene Climatic Optimum on the Dinaride Lake System. Palaeogeography, Palaeoclimatology, Palaeoecology, 299(3), 475–492. 10.1016/j.palaeo.2010.11.024

94. Marek, R. D., & Felice, R. N. (2023). The neck as a keystone structure in avian macroevolution and mosaicism. BMC Biology, 21(1), 216. 10.1186/s12915-023-01715-x

95. Martín-Serra, A., Figueirido, B., Pérez-Claros, J. A., & Palmqvist, P. (2015). Patterns of morphological integration in the appendicular skeleton of mammalian carnivores. Evolution, 69(2), 321–340. 10.1111/evo.12566

96. Martín-Serra, A., Figueirido, B., Pérez-Claros, J. A., & Palmqvist, P. (2015). Patterns of morphological integration in the appendicular skeleton of mammalian carnivores. Evolution, 69(2), 321–340. 10.1111/evo.12566

97. Mayr, G. (2009). Galloanseres. In G. Mayr (Ed.), Paleogene Fossil Birds (pp. 35–59). Springer. 10.1007/978-3-540-89628-9_6

98. McCracken, K. G., Harshman, J., McClellan, D. A., & Afton, A. D. (1999). Data Set Incongruence and Correlated Character Evolution: An Example of Functional Convergence in the Hind-Limbs of Stifftail Diving Ducks. Systematic Biology, 48(4), 683– 714. 10.1080/106351599259979

99. McCurry, M. R., Evans, A. R., Fitzgerald, E. M. G., Adams, J. W., Clausen, P. D., & McHenry, C. R. (2017). The remarkable convergence of skull shape in crocodilians and toothed whales. Proceedings of the Royal Society B: Biological Sciences, 284(1850), 20162348. 10.1098/rspb.2016.2348

100. Michaud, M., Veron, G., & Fabre, A.-C. (2020). Phenotypic integration in feliform carnivores: Covariation patterns and disparity in hypercarnivores versus generalists. Evolution, 74(12), 2681–2702. 10.1111/evo.14112

101. Minh, B. Q., Schmidt, H. A., Chernomor, O., Schrempf, D., Woodhams, M. D., von Haeseler, A., & Lanfear, R. (2020). IQ-TREE 2: New Models and Efficient Methods for Phylogenetic Inference in the Genomic Era. Molecular Biology and Evolution, 37(5), 1530–1534. 10.1093/molbev/msaa015

102. Mlíkovský, J. (2002). Cenozoic birds of the world. Part 1. Europe, 1–406.

103. Mitchell, K. J., Wood, J. R., Scofield, R. P., Llamas, B., & Cooper, A. (2014). Ancient mitochondrial genome reveals unsuspected taxonomic affinity of the extinct Chatham duck (*Pachyanas chathamica*) and resolves divergence times for New Zealand and sub-Antarctic brown teals. Molecular Phylogenetics and Evolution, 70, 420–428. 10.1016/j.ympev.2013.08.017

104. Moen, D. S., Morlon, H., & Wiens, J. J. (2016). Testing Convergence Versus History: Convergence Dominates Phenotypic Evolution for over 150 Million Years in Frogs. Systematic Biology, 65(1), 146–160. 10.1093/sysbio/syv073

105. Morris, Z. S., Vliet, K. A., Abzhanov, A., & Pierce, S. E. (2019). Heterochronic shifts and conserved embryonic shape underlie crocodylian craniofacial disparity and convergence. Proceedings of the Royal Society B: Biological Sciences, 286(1897), 20182389. 10.1098/rspb.2018.2389

106. Natale, R., & Slater, G. J. (2022). The Effects of Foraging Ecology and Allometry on Avian Skull Shape Vary across Levels of Phylogeny. The American Naturalist, 200(4), E174–E188. 10.1086/720745

107. Navalón, G., Bright, J. A., Marugán-Lobón, J., & Rayfield, E. J. (2019). The evolutionary relationship among beak shape, mechanical advantage, and feeding ecology in modern birds*. Evolution, 73(3), 422–435. 10.1111/evo.13655

108. Olsen, A. M. (2015). Exceptional avian herbivores: Multiple transitions toward herbivory in the bird order Anseriformes and its correlation with body mass. Ecology and Evolution, 5(21), 5016–5032. 10.1002/ece3.1787

109. Olsen, A. M. (2017). Feeding ecology is the primary driver of beak shape diversification in waterfowl. Functional Ecology, 31(10), 1985–1995. 10.1111/1365-2435.12890

110. Orkney, A., & Hedrick, B. P. (2024). Small body size is associated with increased evolutionary lability of wing skeleton proportions in birds. Nature Communications, 15(1), 4208. 10.1038/s41467-024-48324-y

111. Orkney, A., Bjarnason, A., Tronrud, B. C., & Benson, R. B. J. (2021). Patterns of skeletal integration in birds reveal that adaptation of element shapes enables coordinated evolution between anatomical modules. Nature Ecology & Evolution, 5(9), Article 9. 10.1038/s41559-021-01509-w

112. Ottenburghs, J. (2016). Crossing Species Boundaries: The Hybrid Histories of the True Geese [Ph.D., Wageningen University and Research]. https://www.proquest.com/docview/2570156611/abstract/B8814004AA9B44FEPQ/1

113. Page, C. E., & Cooper, N. (2017). Morphological convergence in ‘river dolphin’ skulls. PeerJ, 5, e4090. 10.7717/peerj.4090

114. Paradis E, Schliep K (2019). “ape 5.0: an environment for modern phylogenetics and evolutionary analyses in R.” Bioinformatics, 35, 526–528. doi:10.1093/bioinformatics/bty633.

115. Pecsics, T., Laczi, M., Nagy, G., & Csörgő, T. (2017). The cranial morphometrics of the wildfowl (Anatidae). Ornis Hungarica, 25(1), 44–57.

116. Piras, P., Buscalioni, A. D., Teresi, L., Raia, P., Sansalone, G., Kotsakis, T., & Cubo, J. (2014). Morphological integration and functional modularity in the crocodilian skull. Integrative Zoology, 9(4), 498–516. 10.1111/1749-4877.12062

117. Prjibelski, A., Antipov, D., Meleshko, D., Lapidus, A., & Korobeynikov, A. (2020). Using SPAdes De Novo Assembler. Current Protocols in Bioinformatics, 70(1), e102. 10.1002/cpbi.102

118. R Core Team (2024). R: A language and environment for statistical computing. R Foundation for Statistical Computing, Vienna, Austria. URL https://www.R-project.org/.

119. Randau, M., & Goswami, A. (2018). Shape Covariation (or the Lack Thereof) Between Vertebrae and Other Skeletal Traits in Felids: The Whole is Not Always Greater than the Sum of Parts. Evolutionary Biology, 45(2), 196–210. 10.1007/s11692-017-9443-6

120. Randau, M., Cuff, A. R., Hutchinson, J. R., Pierce, S. E., & Goswami, A. (2017). Regional differentiation of felid vertebral column evolution: A study of 3D shape trajectories. Organisms Diversity & Evolution, 17(1), 305–319. 10.1007/s13127-016-0304-4

121. Reding, D. M., Foster, J. T., James, H. F., Pratt, H. D., & Fleischer, R. C. (2008). Convergent evolution of ‘creepers’ in the Hawaiian honeycreeper radiation. Biology Letters, 5(2), 221–224. 10.1098/rsbl.2008.0589

122. Revell, L. J. (2024). phytools 2.0: An updated R ecosystem for phylogenetic comparative methods (and other things). PeerJ, 12, e16505. 10.7717/peerj.16505

123. Rosenblum, E. B., Parent, C. E., & Brandt, E. E. (2014). The Molecular Basis of Phenotypic Convergence. Annual Review of Ecology, Evolution, and Systematics, 45(Volume 45, 2014), 203–226. 10.1146/annurev-ecolsys-120213-091851

124. Salter, J. F., Oliveros, C. H., Hosner, P. A., Manthey, J. D., Robbins, M. B., Moyle, R. G., Brumfield, R. T., & Faircloth, B. C. (2020). Extensive paraphyly in the typical owl family (Strigidae). The Auk, 137(1), ukz070. 10.1093/auk/ukz070

125. Salter, J. F., Hosner, P. A., Tsai, W. L. E., McCormack, J. E., Braun, E. L., Kimball, R. T., Brumfield, R. T., & Faircloth, B. C. (2022). Historical specimens and the limits of subspecies phylogenomics in the New World quails (Odontophoridae). Molecular Phylogenetics and Evolution, 175, 107559. 10.1016/j.ympev.2022.107559

126. Sayyari, E., Whitfield, J. B., & Mirarab, S. (2017). Fragmentary Gene Sequences Negatively Impact Gene Tree and Species Tree Reconstruction. Molecular Biology and Evolution, 34(12), 3279–3291. 10.1093/molbev/msx261

127. Sherratt, E., & Kraatz, B. (2023). Multilevel analysis of integration and disparity in the mammalian skull. Evolution, 77(4), 1006–1018. 10.1093/evolut/qpad020

128. Sherratt, E., & Kraatz, B. (2023). Multilevel analysis of integration and disparity in the mammalian skull. Evolution, 77(4), 1006–1018. 10.1093/evolut/qpad020

129. Sherratt, E., Sanders, K. L., Watson, A., Hutchinson, M. N., Lee, M. S. Y., & Palci, A. (2019). Heterochronic Shifts Mediate Ecomorphological Convergence in Skull Shape of Microcephalic Sea Snakes. Integrative and Comparative Biology, 59(3), 616–624. 10.1093/icb/icz033

130. Sidlauskas, B. (2008). Continuous and arrested morphological diversification in sister clades of characiform fishes: a phylomorphospace approach. Evolution, 62(12), 3135–3156. 10.1111/j.1558-5646.2008.00519.x

131. Song, Y., Wang, Q., An, Z., Qiang, X., Dong, J., Chang, H., Zhang, M., & Guo, X. (2018). Mid-Miocene climatic optimum: Clay mineral evidence from the red clay succession, Longzhong Basin, Northern China. Palaeogeography, Palaeoclimatology, Palaeoecology, 512, 46–55. 10.1016/j.palaeo.2017.10.001

132. Sorenson, M. D., Cooper, A., Paxinos, E. E., Quinn, T. W., James, H. F., Olson, S. L., & Fleischer, R. C. (1999). Relationships of the extinct moa-nalos, flightless Hawaiian waterfowl, based on ancient DNA. Proceedings of the Royal Society of London. Series B: Biological Sciences, 266(1434), 2187–2193. 10.1098/rspb.1999.0907

133. Spencer, L. M. (1995). Morphological Correlates of Dietary Resource Partitioning in the African Bovidae. Journal of Mammalogy, 76(2), 448–471. 10.2307/1382355

134. Sraml, M., Christidis, L., Easteal, S., Horn, P., & Collet, C. (1996). Molecular Relationships Within Australasian Waterfowl (Anseriformes). Australian Journal of Zoology, 44(1), 47–58. 10.1071/zo9960047

135. Stayton, C. T. (2006). Testing Hypotheses of Convergence with Multivariate Data: Morphological and Functional Convergence Among Herbivorous Lizards. Evolution, 60(4), 824–841. 10.1111/j.0014-3820.2006.tb01160.x

136. Stayton, C. T. (2008). Is convergence surprising? An examination of the frequency of convergence in simulated datasets. Journal of Theoretical Biology, 252(1), 1–14. 10.1016/j.jtbi.2008.01.008

137. Stayton, C. T. (2015). The definition, recognition, and interpretation of convergent evolution, and two new measures for quantifying and assessing the significance of convergence. Evolution, 69(8), 2140–2153. 10.1111/evo.12729

138. Steinfield, K. R., Felice, R. N., Kirchner, M. E., & Knapp, A. (2024). Carrion converging: Skull shape predicts feeding ecology in vultures. Journal of Zoology, 322(2), 113–125. 10.1111/jzo.13127

139. Steinthorsdottir, M., Coxall, H. K., de Boer, A. M., Huber, M., Barbolini, N., Bradshaw, C. D., Burls, N. J., Feakins, S. J., Gasson, E., Henderiks, J., Holbourn, A. E., Kiel, S., Kohn, M. J., Knorr, G., Kürschner, W. M., Lear, C. H., Liebrand, D., Lunt, D. J., Mörs, T., … Strömberg, C. a. E. (2021). The Miocene: The Future of the Past. Paleoceanography and Paleoclimatology, 36(4), e2020PA004037. 10.1029/2020PA004037

140. Stidham, T. A., & Zelenkov, N. V. (2017). North American–Asian aquatic bird dispersal in the Miocene: Evidence from a new species of diving duck (Anseriformes: Anatidae) from North America (Nevada) with affinities to Mongolian taxa. Alcheringa: An Australasian Journal of Palaeontology, 41(2), 222–230. 10.1080/03115518.2016.1224439

141. Storer, R. W. (1971). Adaptive radiation of birds. Avian Biology, 1, 149–188.

142. Sun, Z., Pan, T., Hu, C., Sun, L., Ding, H., Wang, H., Zhang, C., Jin, H., Chang, Q., Kan, X., & Zhang, B. (2017). Rapid and recent diversification patterns in Anseriformes birds: Inferred from molecular phylogeny and diversification analyses. PLOS ONE, 12(9), e0184529. 10.1371/journal.pone.0184529

143. Sun, Y., Si, G., Wang, X., Wang, K., & Zhang, Z. (2018). Geometric morphometric analysis of skull shape in the Accipitridae. Zoomorphology, 137(3), 445–456. 10.1007/s00435-018-0406-y

144. Swafford, D. (2003). PAUP*. Phylogenetic Analysis Using parsimony (* and Other Methods). Version 4d10. (No Title). Sinauer Associates, Sunderland, Massachusetts.

145. Tang, R.-X., Wang, J., Li, Y.-F., Zhou, C.-R., Meng, G.-L., Li, F.-J., Lan, Y., Price, M., Podsiadlowski, L., Yu, Y., Wang, X.-M., Liu, Y.-X., Yue, B.-S., Liu, S.-L., Fan, Z.-X., & Liu, S.-Y. (2022). Genomics and morphometrics reveal the adaptive evolution of pikas. Zoological Research, 43(5), 813–826. 10.24272/j.issn.2095-8137.2022.072

146. Tokita, M., Yano, W., James, H. F., & Abzhanov, A. (2017). Cranial shape evolution in adaptive radiations of birds: Comparative morphometrics of Darwin’s finches and Hawaiian honeycreepers. Philosophical Transactions of the Royal Society B: Biological Sciences, 372(1713), 20150481. 10.1098/rstb.2015.0481

147. Tyler, J., & Younger, J. L. (2022). Diving into a dead-end: Asymmetric evolution of diving drives diversity and disparity shifts in waterbirds. Proceedings of the Royal Society B: Biological Sciences, 289(1989), 20222056. 10.1098/rspb.2022.2056

148. Wake, D. B. (1991). Homoplasy: The Result of Natural Selection, or Evidence of Design Limitations? The American Naturalist, 138(3), 543–567. 10.1086/285234

149. Wake, D. B., Wake, M. H., & Specht, C. D. (2011). Homoplasy: From Detecting Pattern to Determining Process and Mechanism of Evolution. Science, 331(6020), 1032– 1035. 10.1126/science.1188545

150. Ward, A. B., Weigl, P. D., & Conroy, R. M. (2002). Functional Morphology of Raptor Hindlimbs: Implications for Resource Partitioning. The Auk, 119(4), 1052–1063. 10.1093/auk/119.4.1052

151. Worthy, T. H. (2009). Descriptions and phylogenetic relationships of two new genera and four new species of Oligo-Miocene waterfowl (Aves: Anatidae) from Australia. Zoological Journal of the Linnean Society, 156(2), 411–454. 10.1111/j.1096-3642.2008.00483.x

152. Worthy, T. H., & Lee, M. S. Y. (2008). Affinities of Miocene Waterfowl (anatidae: Manuherikia, Dunstanetta and Miotadorna) from the St Bathans Fauna, New Zealand. Palaeontology, 51(3), 677–708. 10.1111/j.1475-4983.2008.00778.x

153. Worthy, T. H., Scofield, R. P., Hand, S. J., Pietri, V. L. D., & Archer, M. (2022). <p><strong>A swan-sized fossil anatid (Aves: Anatidae) from the early Miocene St Bathans Fauna of New Zealand</strong></p>. Zootaxa, 5168(1), Article 1. 10.11646/zootaxa.5168.1.3

154. Worthy, T. H., Tennyson, A. J. D., Jones, C., McNamara, J. A., & Douglas, B. J. (2007). Miocene waterfowl and other birds from central Otago, New Zealand. Journal of Systematic Palaeontology, 5(1), 1–39. 10.1017/S1477201906001957

155. Worthy, T. H., Tennyson, A. J. D., Jones, C., McNamara, J. A., & Douglas, B. J. (2007). Miocene waterfowl and other birds from central Otago, New Zealand. Journal of Systematic Palaeontology, 5(1), 1–39. 10.1017/S1477201906001957

156. Zelenkov, N. V. (2020). Cenozoic Evolution of Eurasian Anatids (Aves: Anatidaes. l.). Biology Bulletin Reviews, 10(5), 417–426. 10.1134/S2079086420050096

157. Zelenkov, N. V., & Kurochkin, E. N. (2012). Dabbling ducks (Aves: Anatidae) from the Middle Miocene of Mongolia. Paleontological Journal, 46(4), 421–429. 10.1134/S0031030112040132

158. Zelenkov, N.V. and Martynovich, N.V., Rich fauna of birds of Miocene Tagay site (Olkhon Island, Baikal), Tr. Menzbir. Ornitol. O-va, 2013, no. 2, pp. 73–93.

