## Supplemental Information for "Dietary specialization drives adaptation, convergence, and integration across the cranial and appendicular skeleton in Waterfowl (Anseriformes)"

Appendices

**Appendix I: Phylogenomics of waterfowl relationships**

*UCE recovery* – Sequencing results are summarized in Supplemental Table 1. We obtained an average of 7,938,039 [3,895,081 – 17,901,795] raw reads for toe pad samples, and after trimming, we assembled an average of 217,233 [21,071 – 486,588] contigs. From those contigs, we recovered an average of 3,621 [16 - 4315] UCE loci, however this average is significantly reduced by the small number of UCE loci (16) recovered for *Chauna chavaria*. Despite the small number of loci, *C. chavaria* was consistently recovered as sister to *Chauna torquata* albeit with poor support. We obtained an average of 6,506,220 [5,248 – 16,533,147] raw reads for tissue samples, and after trimming, we assembled an average of 75,811 [4,493 – 163,637] contigs. From those contigs, we recovered an average of 4,215 [239 – 4379] UCE loci. From these data, we produced a 75% taxon complete concatenated matrix of 2,724 loci with 3,819,247 base pairs. Across the matrix, there was a total of 613,630 parsimony informative sites.

**Supplementary Figure 1**. (a) Maximum likelihood topology estimated using IQTree and (b) coalescent topology estimated using SVDquartets implemented in PAUP*. Boot strap values are given at each node. Tip labels include museum accession numbers corresponding to sample voucher used to generate UCE data.

(A)

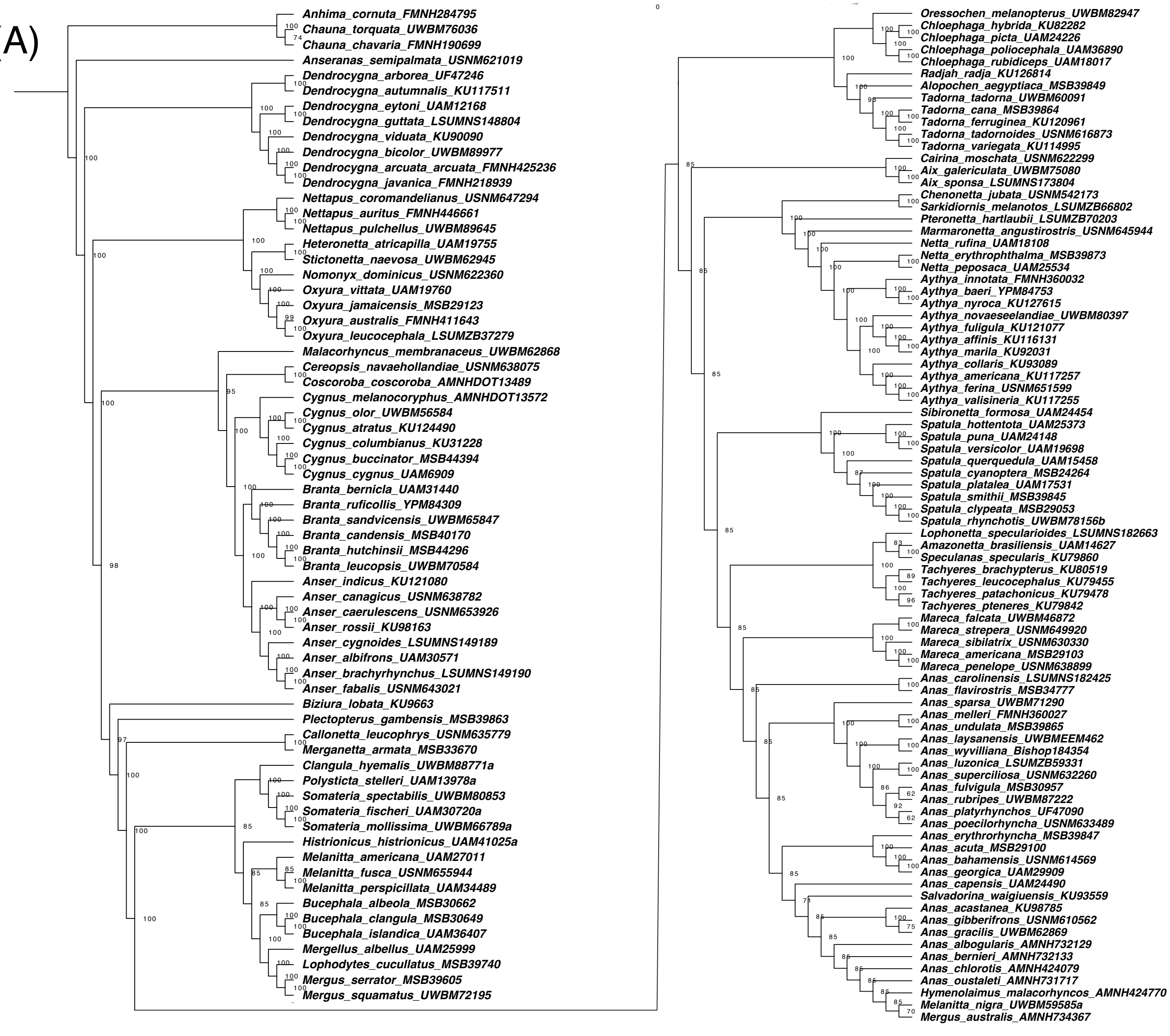

### Appendices

(B)

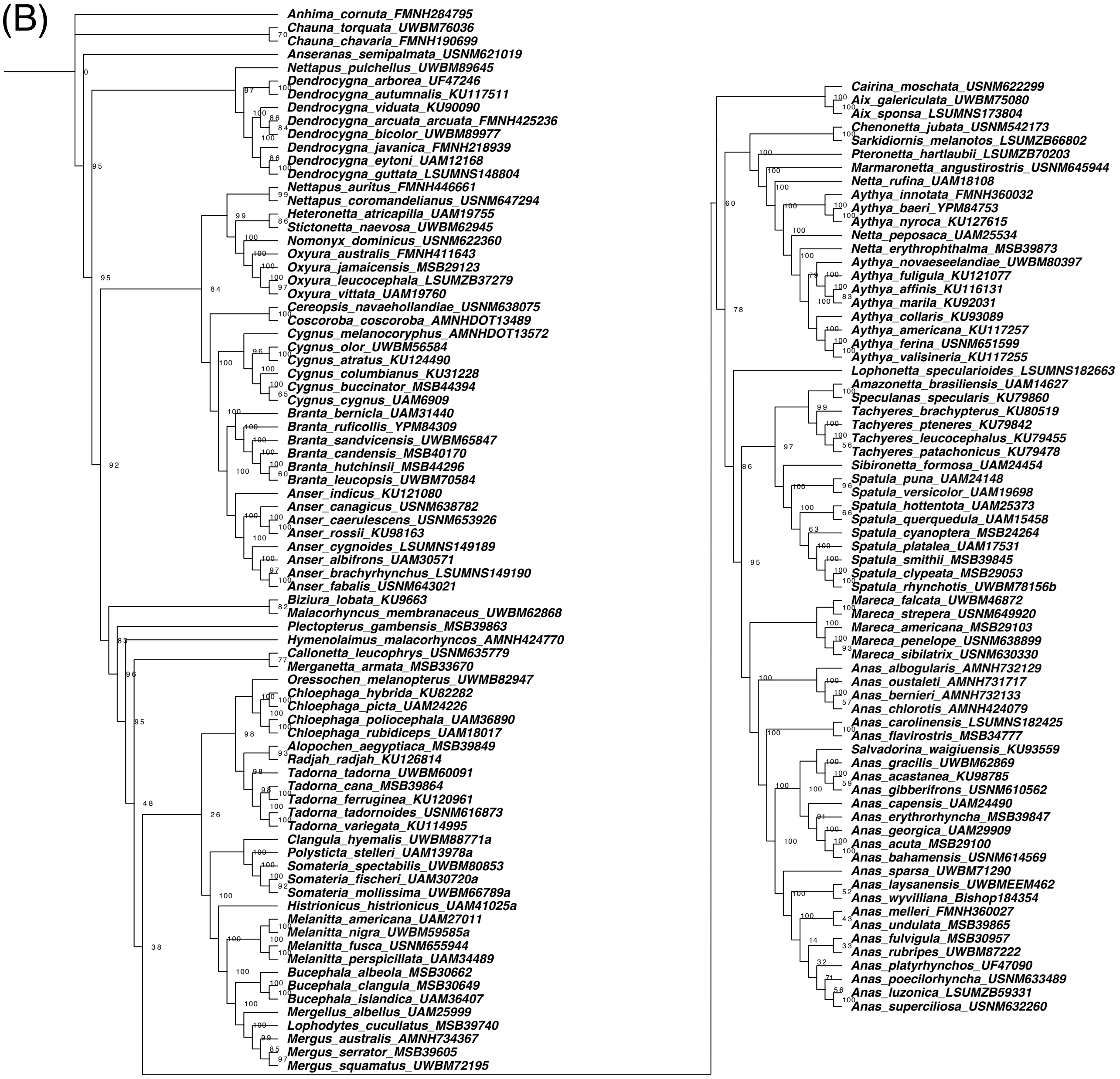

Appendices

**Supplementary Figure 2**. Maximum likelihood topologies estimated using IQTree for "wandering toe pads" belonging to (a) tribe Mergini, (b) genus *Anas*, and (c) *Hymenolaimus*. Tip labels include museum accession numbers corresponding to sample voucher used to generate UCE data.

(A)

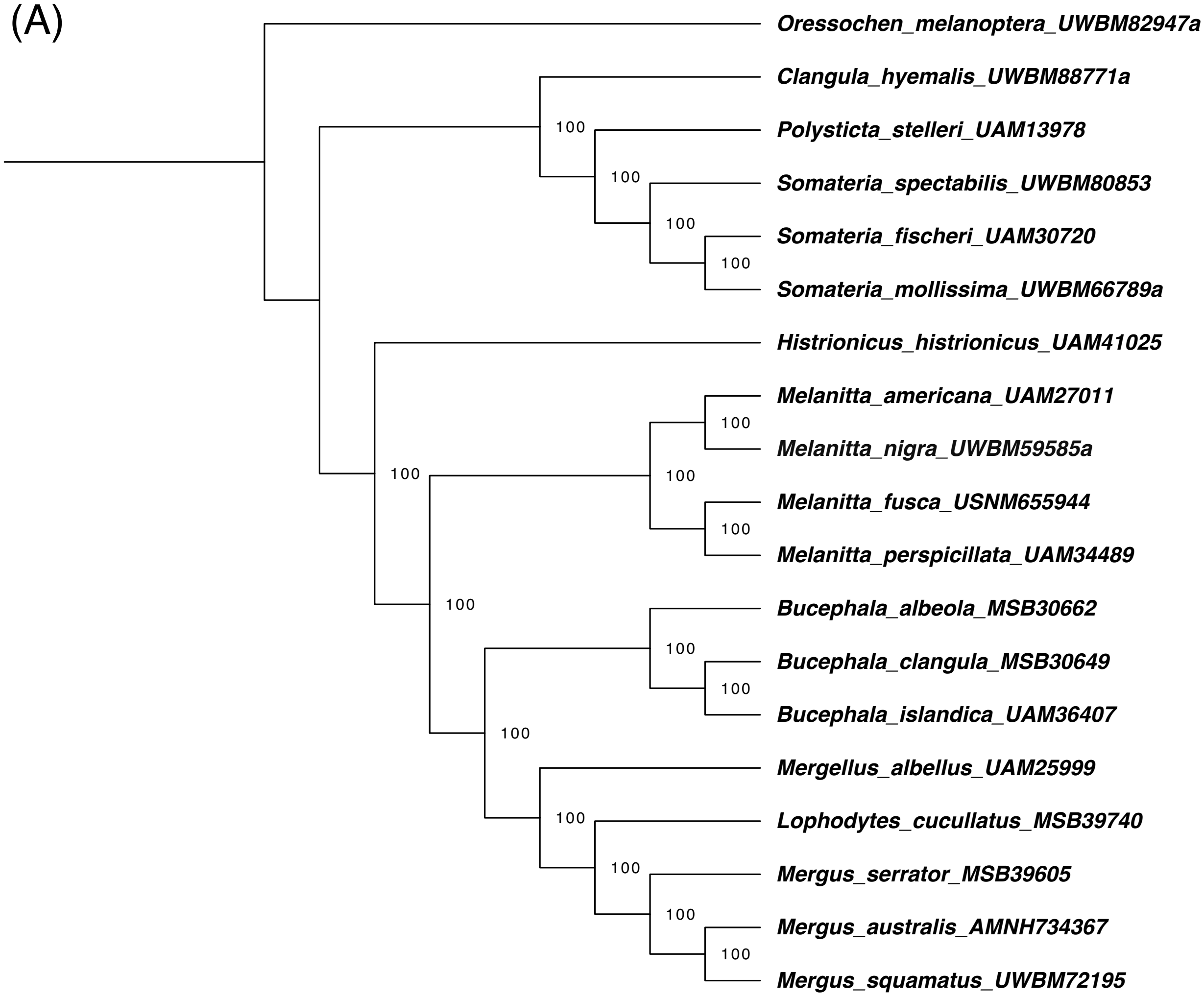

(B)

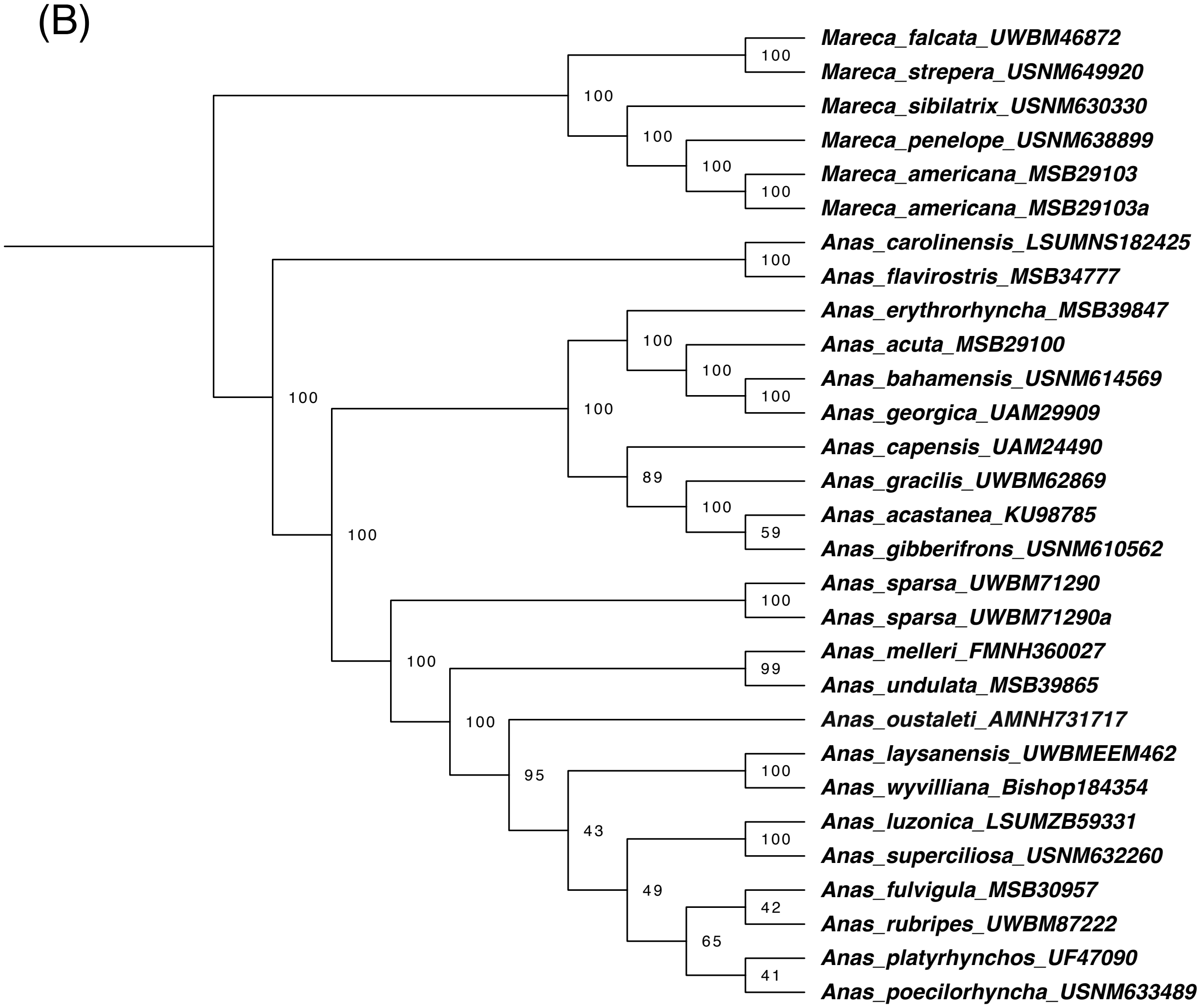
Mareca_falcata_UWBM46872
Mareca_strepera_USNM649920
Mareca_sibilatrix_USNM630330
Mareca_penelope_USNM638899
Mareca_americana_MSB29103
Mareca_americana_MSB29103a
Anas_carolinensis_LSUMNS182425
Anas_flavirostris_MSB34777
Anas_erythrorhyncha_MSB39847
Anas_acuta_MSB29100
Anas_bahamensis_USNM614569
Anas_georgica_UAM29909
Anas_capensis_UAM24490
Anas_gracilis_UWBM62869
Anas_acastanea_KU98785
Anas_gibberifrons_USNM610562
Anas_sparsa_UWBM71290
Anas_sparsa_UWBM71290a
Anas_melleri_FMNH360027
Anas_undulata_MSB39865
Anas_oustaleti_AMNH731717
Anas_laysanensis_UWBMEEM462
Anas_wyvilliana_Bishop184354
Anas_luzonica_LSUMZB59331
Anas_superciliosa_USNM632260
Anas_fulvigula_MSB30957
Anas_rubripes_UWBM87222
Anas_platyrhynchos_UF47090
Anas_poecilorhyncha_USNM633489
100
100
100
100
100
100
100
100
100
100
100
100
89
100
59
100
100
99
100
95
100
43
100
49
42
65
41

Appendices

(C)

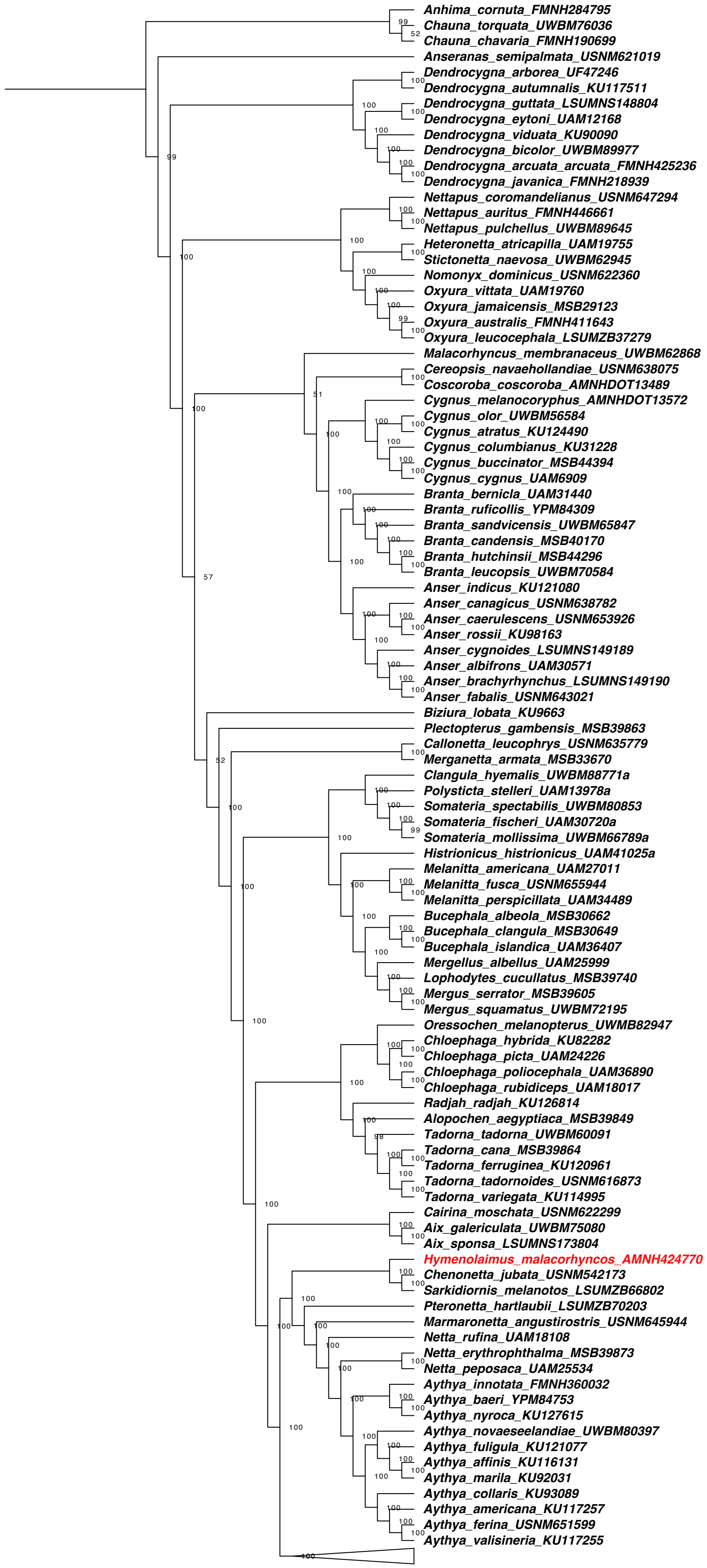

**Supplementary Figure 3**. Maximum likelihood topology estimated using IQTree from a UCE matrix excluding “wandering toe pads”. This topology was fixed and used for the divergence time estimation in BEAST 2. Tip labels include museum accession numbers corresponding to sample voucher used to generate UCE data.

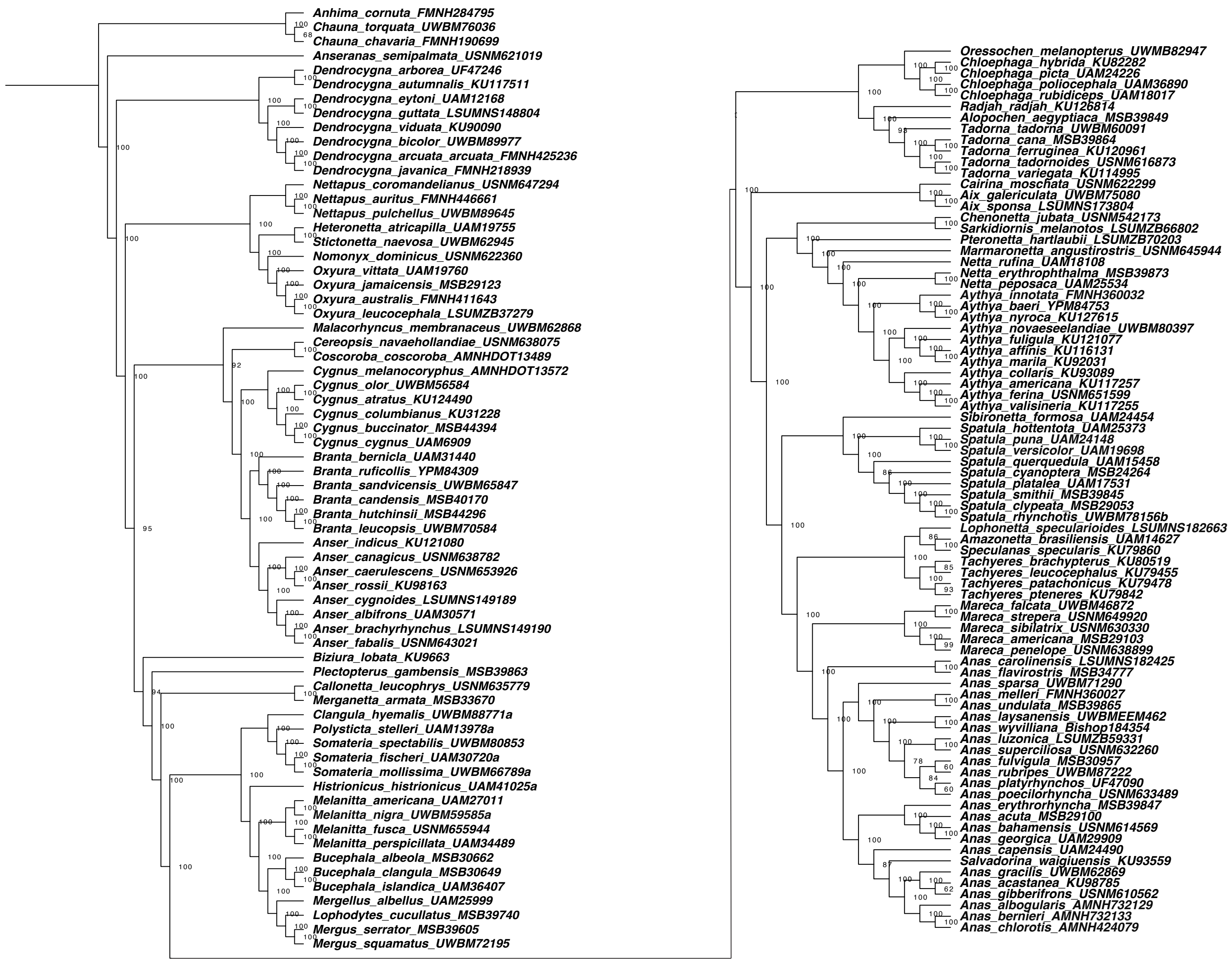

**Appendix II: Landmarking for Geometric Morphometrics**

**Supplementary Figure 4.** Landmarks and semilandmarks used in geometric morphometric analyses. Fixed landmarks are yellow, sliding semi landmarks are shown in red. A, the skull in lateral (i), ventral (ii), dorsal (iii), and caudal view (iv). B Femur in caudal (i) and cranial (ii) view. C Tibiotarsus in anterior (i) and posterior (ii) views. D tarsometatarsus in dorsal (i) and plantar (ii) view.

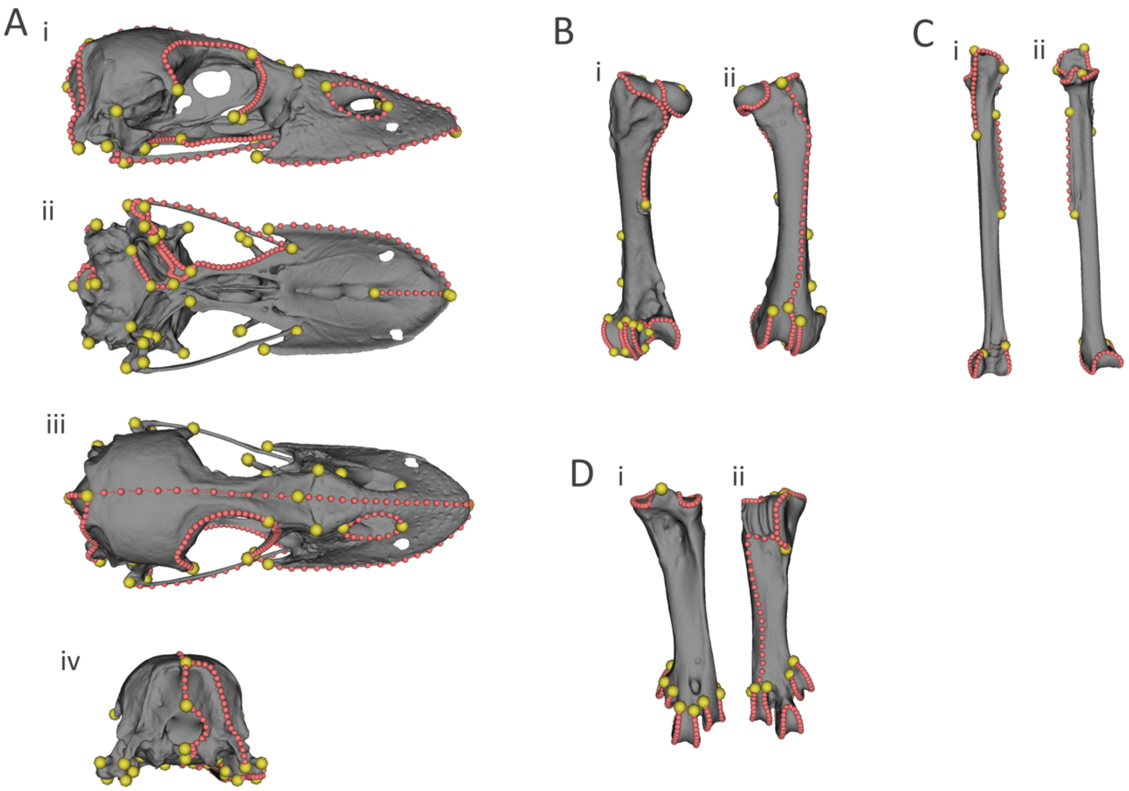

Appendices

Table S3. Landmark numbers and description for the skull.

| Landmark | Location |
|---|---|
| 1 | Tip of Rostrum Palette |
| 2 | Post Choana |
| 3 | R Ventral Jugal Rostrum contact |
| 4 | Anterior Dorsal Tip of Rostrum |
| 5 | Medial Craniofacial hinge |
| 6 | R Posterior Lacrimal Frontal contact orbital margin |
| 7 | R AnteroVentral Pterygoid Palatine contact |
| 8 | R Anterior tip Zygomatic Proccess |
| 9 | Medial Parietal SupraOccipital contact |
| 10 | Medial Dorsal Margin Foramen Magnum |
| 11 | Medial Dorsal Occipital Condyle |
| 12 | Med Ventral Occipital Condyle |
| 13 | Medial Contact BasiOccipital Basisphenoid |
| 14 | Anterior Point Basisphenoid |
| 15 | R PosteroMedial Corner Articular Process Quarate |
| 16 | R AnteriorLateral Articular Process Quarate |
| 17 | R Posterior Pterygoid Quadrate Contact |
| 18 | R Anterior Pterygoid Quadrate Contact |
| 19 | R Anterior Exterior Naris |
| 20 | R Posterior Exterior Naris |
| 21 | R Lateral frontal rostral Contact |
| 22 | L Ventral Jugal Rostral contact |
| 23 | L Post Lacrimal Frontal contact Orbitial margin |
| 24 | L AnterioVentral Pterygoid Palatatine |
| 25 | L Anterior tip Zygomatic Proccess |
| 26 | L PosteroMedial Corner Articular Process Quarate |
| 27 | L AnterioLateral Corner Articular Process Quarate |
| 28 | L Posterior Pterygoid Quadrate Contact |
| 29 | L Anterior Pterygoid Quadrate Contact |
| 30 | L Anterior Exterior Naris |
| 31 | L Posterior Exterior Naris |
| 32 | L Lateral Frontal roastral Contact |
| 33 | R Lateral Posterior Margin Basisphenoid |
| 34 | L Lateral Posterior Margin Basisphenoid |
| 35 | R Anterior Point Lacrimal Ven Margin |
| 36 | R Posterior Point Lacrimal Ven Margin |
| 37 | L Anterior Point Lacrimal Ven Margin |
| 38 | L Posterior Point Lacrimal Ven Margin |
| 39 | R Jugal Quadrate ventral contact |
| 40 | L Jugal Quadrate ventral contact |
| 41 | R Posterior most point of lateral margin of bill |

| | |
|---|---|
| 42 | L Posterior most point of lateral margin of bill |
| 43 | R dorsal most point quadrate |
| 44 | L dorsal most point quadrate |
| 45 | R Ventrolateral most point Supraoccipital |
| 46 | L Ventrolateral most point Supraoccipital |

Table S4. Semi landmark curves for the skull.

| Curve | Location | No. of semilandmarks |
|---|---|---|
| 1 | Anterior Orbit | 25 |
| 2 | Basisphenoid Midline | 5 |
| 3 | Basisphenoid Lateral | 10 |
| 4 | Dorsal Cranium Midline | 13 |
| 5 | Dorsal naris margin | 10 |
| 6 | Dorsal Rostrum | 19 |
| 7 | Foramen Magnum Lateral | 10 |
| 8 | Jugal Bar | 15 |
| 9 | Lateral Rostrum Margin | 19 |
| 10 | Midline Palate | 10 |
| 11 | Occipital complex Lateral | 22 |
| 12 | Occipital Condyle lateral | 7 |
| 13 | Palatine lateral margin | 22 |
| 14 | Pterygoid Lateral Margin | 13 |
| 15 | Pterygoid Midline Margin | 25 |
| 16 | Posterior Orbital Margin | 25 |
| 17 | Quadratocondyle anterior margin | 13 |
| 18 | Quadratocondyle posterior margin | 13 |
| 19 | Supraoccipital midline | 7 |
| 20 | Ventral naris margin | 10 |

### Appendices

Table S5. Landmark numbers and description for the femur

| Landmark number | Location |
|---|---|
| 1 | Posterior turbicle |
| 2 | Crista trochanteris, caudal most point |
| 3 | Distal point of turbicle gastrocnemialis lateralis |
| 4 | Epicondylus lateralis |
| 5 | Medial condyle, proximolateral point on caudal surface |
| 6 | Intercondylar sulcus, distolateral point |
| 7 | Crista tibiofibularis, proximal most point |
| 8 | Lateral condyle, proximal most point of medial margin on cranial surface |
| 9 | Intercondylar sulcus, proximolateral point |
| 10 | Lateral condyle cranial surface depression, proximal most point |
| 11 | Condylar ridge, lateralmost point on caudal surface |
| 12 | Medial condyle cranial surface, proximal most point |
| 13 | Trochlea fibularis, proximocaudal point |
| 14 | Trochlea fibularis, distocaudal point |
| 15 | Trochlea fibularis crista tibiofibularis depression, proximal most point |
| 16 | Trochlea fibularis crista tibiofibularis depression, distalmost point |
| 17 | Proximal point of turbicle gastrocnemialis lateralis |

Table S6. Semi landmark curves for the femur

| Curve | Location | No. of semi landmarks |
|---|---|---|
| 1 | Caudal shaft and latero proximal margin | 40 |
| 2 | Femoral head margin | 34 |
| 3 | Lateral margin of medial condyle and intercondylar sulcus | 9 |
| 4 | Crista tibiofibularis and medial margin of lateral condyle | 40 |
| 5 | Lateral margin of intercondylar sulcus | 28 |
| 6 | Margin of condylar ridge and medial condyle | 37 |
| 7 | Trochlea fibularis | 13 |
| 8 | Trochlea fibularis crista tibiofibularis depression | 13 |
| 9 | Crista trochanteris and cranial shaft | 28 |

Appendices

Table S7. Landmark numbers and description for the tibiotarsus

| Landmark | Location |
|---|---|
| 1 | Cranial cnemial crest, distalmost point |
| 2 | Lateral cnemial crest, lateralmost point |
| 3 | Caudal intercondylar notch |
| 4 | Meeting point of medial margin of proximal articular surface and cranial cnemial crest |
| 5 | Crista fibularis, distalmost point |
| 6 | Crista fibularis, proximal most point |
| 7 | Medial condyle, proximal most point on cranial surface |
| 8 | Lateral condyle, proximal most point on cranial surface |

Table S8. Semi landmark curves for the tibiotarsus

| Curve | Location | No. of semi landmarks |
|---|---|---|
| 1 | Cnemial crests | 27 |
| 2 | Lateral margin of proximal articular surface | 19 |
| 3 | Medial margin of proximal articular surface | 19 |
| 4 | Crista fibularis | 13 |
| 5 | Margins of distal condyles | 39 |

Table S9. Landmark numbers and description for the tarsometatarsus

| Landmark | Location |
|---|---|
| 1 | Eminentia intercotylaris |
| 2 | Crista medialis hypotarsi, dorsoproximal point |
| 3 | Medial crista hypotarsi, dorsodistal point |
| 4 | Dorsoproximal most point medial ridge of trochlea II |
| 5 | Dorsoproximal most point lateral surface of trochlea IV |
| 6 | Trochlea II lateral ridge, proximal most point on dorsal surface |
| 7 | Trochlea II lateral ridge, proximal most point on plantar surface |
| 8 | Trochlea III medial ridge, proximal most point on dorsal surface |
| 9 | Trochlea III lateral ridge, proximal most point on dorsal surface |
| 10 | Trochlea IV medial ridge, proximal most point on its dorsal |
| 11 | Trochlea IV medial ridge, proximal most point on its plantar surface |
| 12 | Trochlea II proximal most point of medial ridge on plantar surface |
| 13 | Trochlea IV proximal most point of lateral ridge on plantar surface |

### Appendices

Table S10. Semi landmark curves for the tarsometatarsus

| Curve | Location | Number of semi landmarks |
|---|---|---|
| 1 | Medial cotyle | 28 |
| 2 | Lateral cotyle | 28 |
| 3 | Crista medialis hypotarsi | 16 |
| 4 | Trochlea II, lateral margin | 22 |
| 5 | Trochlea III margin | 43 |
| 6 | Trochlea IV, medial margin | 22 |
| 7 | Hypotarsi distal margin and plantar shaft | 25 |
| 8 | Trochlea II, medial margin | 19 |
| 9 | Trochlea IV lateral margin | 19 |

Table S12. Species added to the tree modified for use in morphometric analyses

| Species | Divergence date (ma) | Source |
|---|---|---|
| *Hymenolaimus malacorhynchus* | 14 | Fulton et al. 2012 |
| *Cyanochen cyanopterus* | 7.3 | Sun et al. 2017 |
| *Anas aucklandica* | 1 | Mitchell et al. 2014 |
| *Anser anser* | 0.9 | Sun et al. 2017 |
| *Neochen jubata* | 6.6 | Sun et al. 2017 |
| *Oxyura maccoa* | 4.2 | Sun et al. 2017 |
| *Asarcornis scutulata* | 10 | Fulton et al. 2012 |

Table 13. Pairwise probabilities of net evolutionary rate differences for the skull based on ecotype. Numbers in bold indicate a significant relationship.

| | Dive_grasp/Diving | Fish/Diving | Herbivore/Surface | Herbivore/Terrestrial | Herbivore/Wading | Mixed/Surface | Mixed/Wading | Straining/Diving |
|---|---|---|---|---|---|---|---|---|
| Fish/Diving | **0.002** | | | | | | | |
| Herbivore/Surface | **0.002** | **0.001** | | | | | | |
| Herbivore/Terrestrial | 0.052 | **0.017** | **0.001** | | | | | |
| Herbivore/Wading | **0.018** | **0.001** | 0.961 | **0.001** | | | | |
| Mixed/Surface | 0.447 | **0.001** | **0.009** | **0.007** | 0.052 | | | |
| Mixed/Wading | **0.001** | **0.001** | **0.001** | **0.001** | **0.001** | **0.001** | | |
| Straining/Diving | **0.001** | **0.001** | 0.441 | **0.001** | 0.625 | **0.001** | **0.001** | |
| Straining/Surface | **0.034** | **0.001** | **0.056** | **0.001** | 0.186 | 0.158 | **0.001** | **0.002** |

Appendices

Table 14. Pairwise probabilities of net evolutionary rate differences for the femur, tibiotarsus, and tarsometatarsus based on foraging type. Numbers in bold indicate a significant relationship.

| | | | |
|---|---|---|---|
| **Femur** | | | |
| | Diving | Surface | Terrestrial |
| Surface | **0.017** | | |
| Terrestrial | **0.001** | **0.001** | |
| Wading | **0.001** | **0.001** | **0.001** |
| **Tibiotarsus** | | | |
| | Diving | Surface | Terrestrial |
| Surface | **0.001** | | |
| Terrestrial | 0.428 | **0.001** | |
| Wading | **0.001** | **0.001** | **0.001** |
| **Tarsometatarsus** | | | |
| | Diving | Surface | Terrestrial |
| Surface | 0.339 | | |
| Terrestrial | 0.111 | **0.013** | |
| Wading | **0.001** | **0.001** | **0.001** |

Appendices

**Appendix III: Multi-locus analysis of waterfowl relationships**

For comparison to the UCE analysis, we present here a previously unpublished genus-level phylogeny for the waterfowl based on portions of four nuclear genes and the mitochondrial small subunit (12S) rRNA gene (Supplementary Table X1). The data set includes 84 taxa (Supplementary Table X2), including representatives of all extant waterfowl genera recognized by Clements et al. (2023), including the likely extinct pink-headed duck (*Rhodonessa caryophyllacea*) but not the extinct Labrador duck (*Camptorhynchus labradorius*). A few species were included in the analysis based on 12S data only (see Supplementary Table X2). Most of the samples used for this analysis were assembled in the mid-1990's, relying heavily on feather samples obtained from: 1) live birds in the captive waterfowl collection at what is now the Sylvan Heights Bird Park in North Carolina; and 2) the Migratory Bird Parts Collection Survey coordinated by the United States Fish & Wildlife Service. Additional samples were provided by museum collections or colleagues (Supplementary Table X2). Most of the DNA sequence data were generated between 1993 and 2006, and approximately half of the sequences were submitted to GenBank in conjunction with earlier publications (Bulgarella et al. 2010; García-Moreno et al. 2003; McCracken et al. 2009; McCracken & Sorenson 2005; Mindell et al. 1997; Sorenson et al. 2003). Here, we assemble and augment the previously published data to yield a more comprehensive analysis with a complete sampling of extant waterfowl genera.

DNA extraction, PCR, DNA sequencing, reconciling of forward and reverse reads, and sequence alignment followed standard methods, as described in the references cited above. PCR and sequencing primers, including internal primers to amplify smaller fragments from museum specimens (e.g., *Rhodonessa*) are listed in Supplementary Table X*3*. Data for all four nuclear genes plus the mitochondrial 12S gene were combined into a single data set and analyzed using equal-weights parsimony in PAUP* (Swofford, 2002). For the nuclear genes, multiple-base insertions and deletions (indels) were common, so each unique indel, regardless of length, was recoded as a single binary character indicating presence or absence of the indel. A set of binary indel characters for each nuclear gene was added to the data matrix and gap characters in the sequence alignment were replaced with question marks (signifying missing data). Examination of the 12S alignment indicates that a strong preponderance of indels involved insertions or deletion of a single base, so gaps were treated as a fifth character state ("gapmode=newstate") for 12S only. To thoroughly explore tree space and to find equally parsimonious "islands" of trees, we completed 1000 replicate searches with random addition of taxa. We also completed a bootstrap analysis based on 1000 resampled data sets, each of which was analyzed in a similar manner (i.e., with multiple searches and random addition of taxa). The NEXUS file analyzed in PAUP*, including the commands used, is available as a supplementary document.

We found sixteen equally parsimonious trees of length 6,317 in two islands of eight trees, one of which is illustrated in Supplementary Fig. X1. The 16 trees differed only in: 1) species-level relationships within *Oxyura*; 2) species-level relationships within the dabbling ducks (tribe

Anatini); and 3) the positions of two relatively divergent, monotypic lineages, ringed teal *Callonetta* and torrent duck *Merganetta*. Overall, this data set supports several key results reported in other molecular analyses of the waterfowl (e.g., Johnson & Sorenson 1999; Donne-Goussé et al. 2002; McCracken & Sorenson 2005; Gonzalez et al. 2009; Bulgarella et al. 2010; Sun et al. 2017; Buckner et al. 2018), all of which involve relationships that were not evident in morphological analyses of the clade (Livezey 1997). These include: 1) the sister relationship of pygmy geese (*Nettapus* spp.) and stifftails (traditionally assigned to tribe Oxyurini) and the divergence of this clade from other "ducks" (subfamily Anatidae); 2) the inclusion of freckled duck *Stictonetta* (traditionally viewed as a transitional form between the Anserine and Anatinae) in the stifftail clade; 3) the divergence of musk duck *Biziura* and pink-eared duck *Malacorhynchus*, respectively, from all other waterfowl; 4) the sister relationship between *Coscoroba* and *Cereopsis*; 5) the divergence of spur-winged goose *Plectropterus*, ringed teal *Callonetta* and torrent duck *Merganetta*, respectively, from other "ducks" (i.e., within the subfamily Anatinae); 6) the sister relationship between *Pteronetta* and *Cyanochen*; and 7) the close relationship of *Asarcornis* to the pochards (tribe Aythini). In addition, the placement of Salvadori's teal (*Anas* "*Salvadorina*" *waigiuensis*) within the genus *Anas*, a result also recovered in the UCE analysis, has to our knowledge not been reported in previous molecular studies. Overall, the results of this multi-locus analysis accord well with the results of the UCE analysis, with almost all well-supported nodes in the former tree present in the latter. The UCE analysis, however, helps to resolve many of the poorly resolved relationships in the multi-locus analysis. This includes the relationships among subfamily-level lineages within Anatidae and the relationships among tribes within the "ducks" (subfamily Anatinae). As enumerated in Supplementary Figure 5, we suggest that there are six or seven comparably divergent lineages that should be recognized as anatid subfamilies, and eight to ten comparably divergent lineages that should be recognized as tribes within the subfamily Anatinae. Both lists include a larger number of lineages than have been included recent taxonomic treatments of the waterfowl (e.g., Livezey 1997; Winkler et al. 2015) but all the root names we suggest have been used to refer to a proposed waterfowl subfamily in the historical literature (Bock 1994). Note that the composition of several "traditional" tribes (e.g., Anatini, Aythyini, Tadornini, Cairinini, Oxyurini) differs from earlier taxonomic treatments; ours and previous molecular analyses provide a much-improved understanding of waterfowl relationships.

**Supplementary Figure 5.** One of 16 equally parsimonious trees with bootstrap values indicated at each node (with values <70 highlighted in red). Genera not included in the UCE data set are highlighted in red. Suggested waterfowl subfamilies and tribes (within Anatinae) are listed with the corresponding lineages in the tree indicated with black and red numbering, respectively. The ordering of subfamilies and tribes in the numbered lists reflect the results of the UCE analysis.

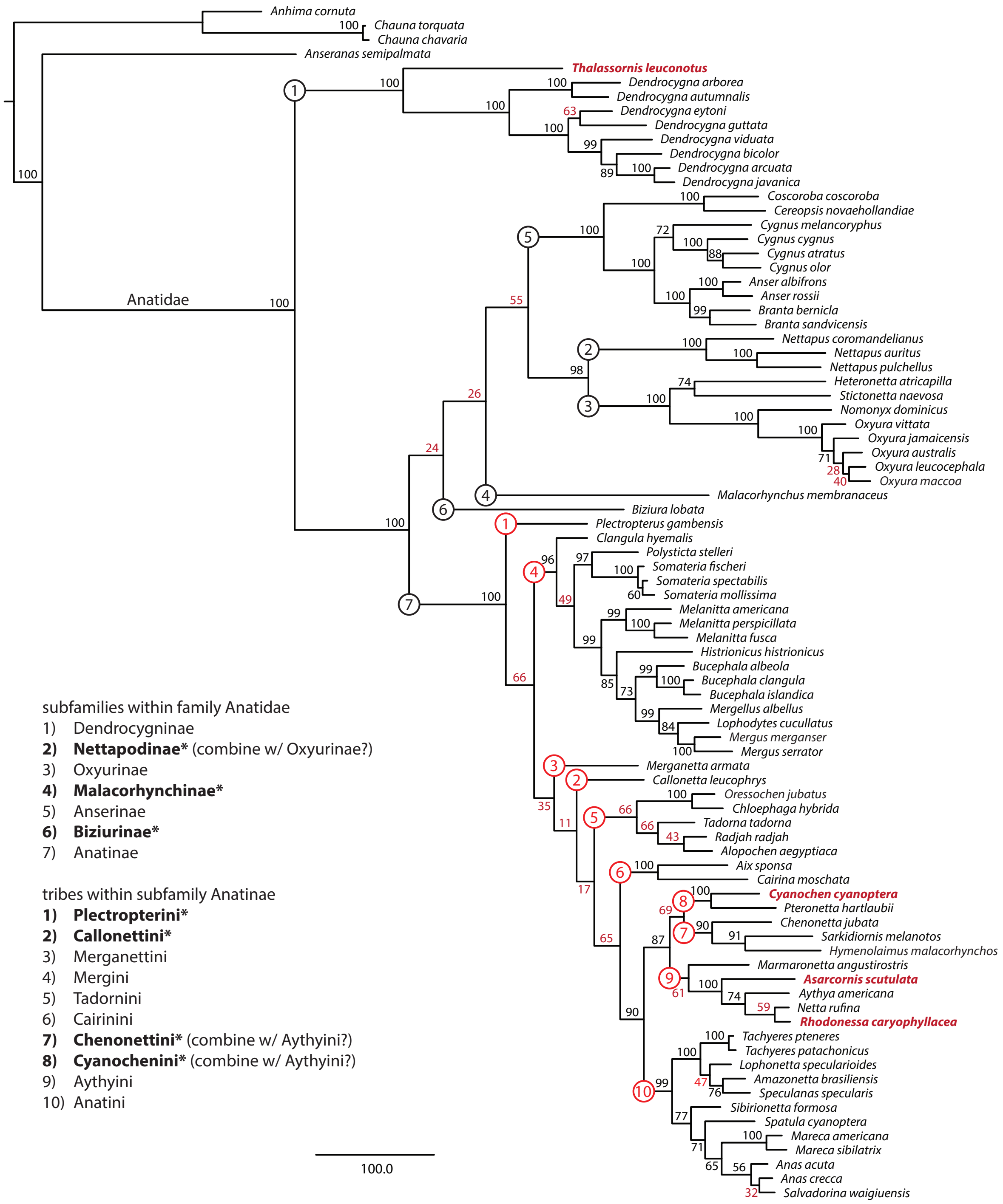

**Supplementary Table X1.** Gene regions included in multi-locus analysis of waterfowl relationships.

| Gene | Location in mallard reference genome | Length (base pairs) coding : non-coding | Previously published* sequences (reference: GenBank accession #s) |
|---|---|---|---|
| CD4 membrane glycoprotein of T lymphocytes (CD4) – intron 4 | NC_051772.1: 82,214,614-82,214,193 | 77 : 345-445 | 1: HM063614-HM063640 |
| hemoglobin subunit alpha-A – coding regions and introns (H-alpha-A) | NC_051786.1: 6,824,524-6,823,826 | 429 : 255-298 | 3: GQ271019-GQ271711 |
| lecithin cholesterol acyltransferase (LCAT) – introns 2-4 | NC_051783.1: 12,065,227-12,063,907 | 434 : 876-1156 | 1: HM063584-HM063611 |
| phosphoenolpyruvate carboxykinase, cytosolic (PCK1) – intron 3 | NC_051792.1: 9,157,497-9,156,850 | 17 : 598-620 | 1: HM063643-HM063670<br>4: AY747841-AY747845 |
| phosphoenolpyruvate carboxykinase, cytosolic (PCK1) – intron 9 | NC_051792.1: 9,153,608-9,152,923 | 92 : 581-598 | 1: HM063673-HM063699<br>4: AY747846-AY747851<br>6: AY274082-AY274112 |
| mitochondrial small subunit ribosomal RNA (12S) plus portions of flanking tRNA genes | NC_009684.1: 1,085-2,112 | 0 : 1014-1044 | 1: HM063531-HM063558<br>2: AF536740<br>4: AY747698-AY747703<br>5: U83728-U83739<br>6: AY274006 |

* references: 1) Bulgarella et al. (2010); 2) García-Moreno et al. (2003); 3) McCracken et al. (2009); 4) McCracken & Sorenson (2005); 5) Mindell et al. (1997); 6) Sorenson et al. (2003).

**Supplementary Table X2.** List of samples and GenBank accession numbers for the "multi-locus" data set. Taxonomy follows Clements et al. (2023). Acronyms for sample sources include: LSUNMS: Louisiana State Museum of Natural Science; MMNH: University of Minnesota Bell Museum of Natural History; SHBP: Sylvan Heights Bird Park; UAM: University of Alaska Museum of the North; UMMZ: University of Michigan Museum of Zoology; UMN: University of Minnesota; USFWS: United States Fish & Wildlife Service Migratory Bird Parts Collection Survey; USNM: United States National Museum of Natural History; USNZP: United States National Zoological Park; UWBM: University of Washington Burke Museum of Natural History and Culture.

| | | GenBank Accession Numbers | | | | | | |
|---|---|---|---|---|---|---|---|---|
| **Species** | **Sample ID** | **CD4** | **H-alpha-a** | **LCAT** | **PCK1:3** | **PCK1:9** | **12S** | **Source: Geographic origin** |
| *Anhima cornuta* | ANHI | — | — | — | TBD | — | U83728* | UMMZ 152361: captive, Detroit Zoo |
| *Chauna torquata* | CHTO | TBD | GQ271467* | TBD | TBD | AY274082 | AY274006 | UMMZ 235046 (T1331): captive, Detroit Zoo |
| *Chauna chavaria* | CHCH | — | — | — | — | — | U83729* | UMMZ 150868: captive, Detroit Zoo |
| *Anseranas semipalmata* | ANSE | TBD | GQ271466* | TBD | TBD | AY274083 | U83730 | SHBP: captive |
| *Thalassornis leuconotus* | THLE | TBD | GQ271468* | TBD | TBD | TBD | U83739* | SHBP: captive |
| *Dendrocygna arborea* | DEAB | TBD | GQ271473* | TBD | TBD | TBD | TBD | N.L. Staus, U. Minnesota: Hog Cay, Long Island, Bahamas |
| *Dendrocygna autumnalis* | DEAU | TBD | GQ271471* | TBD | TBD | TBD | TBD | USFWS: Brevard Co., FL |
| *Dendrocygna viduata* | DEVI | TBD | GQ271470* | TBD | TBD | TBD | TBD | SHBP: captive |
| *Dendrocygna bicolor* | DEBI | TBD | GQ271474* | TBD | TBD | TBD | U83737* | USFWS: Brevard Co., FL |
| *Dendrocygna guttata* | DEGU | TBD | GQ271472* | TBD | TBD | TBD | TBD | SHBP: captive |
| *Dendrocygna eytoni* | DEEY | TBD | GQ271476* | TBD | TBD | TBD | TBD | SHBP: captive |
| *Dendrocygna arcuata* | DEAR | TBD | GQ271477* | TBD | TBD | AY274084 | AF536740 | Musem Victoria (MV1923): Finley, NSW |
| *Dendrocygna javanica* | DEJA | TBD | GQ271478* | TBD | TBD | TBD | TBD | SHBP: captive |
| *Coscoroba coscoroba* | COCO | TBD | GQ271483* | TBD | TBD | TBD | TBD | SHBP: captive |
| *Cereopsis novaehollandiae* | CENO | TBD | GQ271485* | TBD | TBD | TBD | TBD | SHBP: captive |
| *Cygnus melancoryphus* | CYME | TBD | GQ271488* | TBD | TBD | TBD | TBD | SHBP: captive |
| *Cygnus cygnus* | CYCY | TBD | GQ271495* | TBD | TBD | TBD | TBD | SHBP: captive |
| *Cygnus atratus* | CYAT | TBD | GQ271490* | TBD | TBD | TBD | U83731* | SHBP: captive |
| *Cygnus olor* | CYOL | HM063640 | GQ271492* | HM063611 | HM063670 | HM063699 | HM063558 | SHBP: captive |
| *Branta bernicla* | BRBE | HM063639 | GQ271510* | HM063610 | HM063669 | HM063698 | HM063557 | MMNH 38648 (KSW505): Cape Pierce, AK |
| *Branta sandvicensis* | BRSA | TBD | GQ271509* | TBD | TBD | TBD | U83735* | SHBP: captive |
| *Anser albifrons* | ANAL | TBD | GQ271501* | TBD | TBD | TBD | TBD | USFWS: Cameron Co., LA |
| *Anser rossii* | ANRO | TBD | GQ271506* | TBD | TBD | TBD | U83734* | USFWS: Burke Co., ND |
| *Nettapus coromandelianus* | NECO | TBD | GQ271517* | TBD | TBD | TBD | TBD | SHBP: captive |
| *Nettapus auritus* | NEAU | TBD | GQ271516* | TBD | TBD | TBD | TBD | SHBP: captive |
| *Nettapus pulchellus* | NEPU | TBD | GQ271518* | TBD | TBD | TBD | TBD | SHBP: captive |
| *Heteronetta atricapilla* | HEAT | TBD | GQ271522* | TBD | TBD | TBD | TBD | SHBP: captive |

| | | | | | | | | |
|---|---|---|---|---|---|---|---|---|
| *Stictonetta naevosa* | STNA | TBD | GQ271520* | TBD | TBD | TBD | TBD | SHBP: captive |
| *Nomonyx dominicus (1)* | NODO | TBD | GQ271524* | TBD | TBD | AY747846 | — | UAM 14784 (KGM464): Corrientes, Argentina |
| *Nomonyx dominicus (2)* | NODO | — | — | — | — | — | AY747698 | LSUNMS 123431: Santa Cruz, Bolivia |
| *Oxyura vittata* | OXVI | TBD | GQ271526* | TBD | AY747841 | AY747847 | AY747699 | SHBP: captive |
| *Oxyura jamaicensis* | OXJA | TBD | GQ271528* | TBD | AY747842 | AY747848 | AY747700 | USFWS: Siskiyou Co., CA |
| *Oxyura australis* | OXAU | TBD | GQ271532* | TBD | AY747843 | AY747849 | AY747701 | Musem Victoria (MV1991): Australia |
| *Oxyura leucocephala* | OXLE | TBD | GQ271530* | TBD | AY747844 | AY747850 | AY747702 | SHBP: captive |
| *Oxyura maccoa* | OXMA | TBD | GQ271534* | TBD | AY747845 | AY747851 | AY747703 | SHBP: captive |
| *Biziura lobata* | BILO | TBD | GQ271480* | TBD | TBD | TBD | TBD | Musem Victoria (MV785): Australia |
| *Malacorhynchus membranaceus* | MAME | TBD | GQ271515* | TBD | TBD | TBD | TBD | SHBP: captive |
| *Callonetta leucophrys* | CALE | HM063629 | GQ271550* | HM063599 | HM063658 | HM063688 | HM063546 | UMN (Cedar Creek Natural History Area): captive |
| *Plectropterus gambensis* | PLGA | TBD | GQ271535* | TBD | TBD | TBD | TBD | SHBP: captive |
| *Merganetta armata* | MEAR | HM063638 | GQ271063* | HM063609 | HM063668 | HM063697 | HM063556 | UWBM 54417: Tucumán, Argentina |
| *Cairina moschata* | CAMO | HM063625 | GQ271544* | HM063595 | HM063654 | HM063684 | HM063542 | SHBP: captive |
| *Aix sponsa* | AISP | HM063626 | GQ271545* | HM063596 | HM063655 | HM063685 | HM063543 | SHBP: captive |
| *Oressochen jubatus* | NEJU | HM063636 | GQ271019* | HM063607 | HM063666 | HM063696 | HM063554 | SHBP: captive |
| *Chloephaga hybrida* | CHHY | TBD | GQ271024* | TBD | TBD | TBD | TBD | MMNH 42140 (JK93148): Ushuaia Bay, Tierra del Fuego, Argentina |
| *Tadorna tadorna* | TATA | HM063627 | GQ271540* | HM063597 | HM063656 | HM063686 | HM063544 | SHBP: captive |
| Radjah radjah | TARA | — | — | — | — | — | TBD | SHBP: captive |
| *Alopochen aegyptiaca* | ALAE | HM063635 | GQ271538* | HM063606 | HM063665 | HM063695 | HM063553 | SHBP: captive |
| *Clangula hyemalis* | CLHY | TBD | GQ271593* | TBD | TBD | TBD | TBD | USFWS: Hancock Co., ME |
| *Polysticta stelleri* | POST | TBD | GQ271573* | TBD | TBD | TBD | TBD | USFWS: Kenai, AK |
| *Somateria fischeri* | SOFI | TBD | GQ271575* | TBD | TBD | TBD | U83738* | UWBM 44257: Yakutiya, Russia |
| *Somateria mollissima* | SOMO | TBD | GQ271583* | TBD | TBD | TBD | TBD | USFWS: Hancock Co., ME |
| *Somateria spectabilis* | SOSP | TBD | GQ271577* | TBD | TBD | TBD | TBD | MMNH 39007 (JKAK9066) : Chagvan Bay, Togiak NWR, AK |
| *Histrionicus histrionicus* | HIHI | TBD | GQ271585* | TBD | TBD | TBD | TBD | USFWS: Kenai, AK |
| *Bucephala clangula* | BUCL | TBD | GQ271597* | TBD | TBD | TBD | TBD | USFWS: Thurston Co., CA |
| *Bucephala islandica* | BUIS | TBD | GQ271598* | TBD | TBD | TBD | TBD | USFWS: Park Co., MT |
| *Bucephala albeola* | BUAL | TBD | GQ271595* | TBD | TBD | TBD | TBD | USFWS: Douglas Co., WA |
| *Melanitta americana* | MENI | TBD | GQ271591* | TBD | TBD | TBD | TBD | USFWS: Carteret Co., NC |
| *Melanitta perspicillata* | MEPE | TBD | GQ271587* | TBD | TBD | TBD | TBD | MMNH 38661 (KSW512) : Cape Pierce, AK |
| *Melanitta fusca* | MEFU | TBD | GQ271589* | TBD | TBD | TBD | TBD | USFWS: Hancock Co., ME |
| *Mergellus albellus* | MEAL | TBD | GQ271600* | TBD | TBD | TBD | TBD | USNZP (213647): captive |
| *Lophodytes cucullatus* | MECU | TBD | GQ271602* | TBD | TBD | TBD | TBD | USFWS: Yuba Co., CA |

| | | | | | | | | |
|---|---|---|---|---|---|---|---|---|
| *Mergus merganser* | MEME | TBD | GQ271604* | TBD | TBD | TBD | TBD | USFWS: Illinois |
| *Mergus serrator* | MESE | TBD | GQ271606* | TBD | TBD | TBD | TBD | USFWS: Brazoria Co., TX |
| *Pteronetta hartlaubii* | PTHA | HM063632 | GQ271537* | HM063602 | HM063661 | HM063691 | HM063549 | SHBP: captive |
| *Cyanochen cyanoptera* | CYCN | HM063633 | GQ271051* | HM063603 | HM063662 | HM063692 | HM063550 | SHBP: captive |
| *Sarkidiornis melanotos* | SAME | HM063624 | GQ271536* | HM063594 | HM063653 | HM063683 | HM063541 | SHBP: captive |
| *Hymenolaimus malacorhynchos* | HYMA | TBD | TBD | HM063605 | HM063664 | HM063694 | HM063552 | DNA extract via M. Williams and A. Cooper: Manganuiateao River, NZ |
| *Chenonetta jubata* | CHJU | HM063628 | GQ271548* | HM063598 | HM063657 | HM063687 | HM063545 | SHBP: captive |
| *Asarcornis scutulata* | CASC | HM063631 | GQ271552* | HM063601 | HM063660 | HM063690 | HM063548 | SHBP: captive |
| *Netta rufina* | NERU | TBD | GQ271553* | TBD | TBD | TBD | TBD | SHBP: captive |
| *Aythya americana* | AYAM | HM063630 | GQ271558* | HM063600 | HM063659 | AY274112 | HM063547 | M.C. Woodin, Texas A&M (LMTX R654): Laguna Madre, TX |
| *Marmaronetta angustirostris* | MAAN | HM063634 | GQ271551* | HM063604 | HM063663 | HM063693 | HM063551 | SHBP: captive |
| *Tachyeres patachonicus* | TAPA | HM063614 | GQ271608* | HM063584 | HM063643 | HM063673 | HM063531 | SHBP: Chile, museum specimen |
| *Tachyeres pteneres* | TAPT | HM063615 | GQ271613* | HM063585 | HM063644 | HM063674 | HM063532 | DNA extract via G.B. Nunn and E.E. Paxinos: Navarino Is., Chile |
| *Amazonetta brasiliensis* | AMBR | HM063616 | GQ271619* | HM063586 | HM063645 | HM063675 | HM063533 | SHBP: captive |
| *Speculanas specularis* | ANSP | HM063617 | GQ271616* | HM063587 | HM063646 | HM063676 | HM063534 | SHBP: captive |
| *Lophonetta specularioides* | LOSP | HM063618 | GQ271064* | HM063588 | HM063647 | HM063677 | HM063535 | SHBP: captive |
| *Spatula cyanoptera* | ANCY | HM063619 | GQ271145* | TBD | TBD | TBD | TBD | MMNH 42277 (JK95137): nw of Buno Aires, Argentina |
| *Sibirionetta formosa* | ANFO | TBD | GQ271703* | TBD | TBD | TBD | U83732* | MMNH 42241 Cedar Creek (X7134): captive |
| *Mareca sibilatrix* | MASI | TBD | GQ271710* | TBD | TBD | TBD | TBD | LSUMNS B-20764: captive |
| *Mareca americana* | MAAM | HM063620 | GQ271711* | HM063590 | HM063649 | HM063679 | HM063537 | USFWS: Brazoria Co., TX |
| *Anas acuta (1)* | ANAC | HM063621 | GQ271714 | — | — | HM063680 | — | UWBM 43948: Russia |
| *Anas acuta (2)* | ANAC | — | — | HM063591 | HM063650 | — | HM063538 | USNZP: captive |
| *Anas crecca (1)* | ANCR | HM063623 | — | HM063593 | HM063652 | HM063682 | HM063540 | SHBP: captive |
| *Anas crecca (2)* | ANCR | — | GQ271739 | — | — | — | — | UAM 9597 (KSW 2960): Adak Island, AK |
| *Salvadorina waigiuensis* | SAWA | — | — | — | — | — | TBD | USNM 518907: museum specimen, Irian Jaya, Wamena, Baliem Valley |
| *Rhodonessa caryophyllacea* | RHCA | — | — | — | — | — | TBD | DNA extract via USNM and E.E. Paxinos: museum specimen |

**Supplementary Table X3.** Primers used to amplify and sequence gene regions in Supplementary Table X1. Primers used for most high-quality samples are indicated with an asterisk; additional primers were used to amplify smaller fragments in lower-quality samples and/or to facilitate nested PCR protocols.

| Primers for nuclear loci | Primers for 12S rRNA |
|---|---|
| *CD4.4F CTCCATCGATTAATNAGAACATCTCC<br>*CD4.5R TTCCKGAAGTTCAGAYGCCATGAC<br>*HaA.F1 GGGCACCCGTGCTGGGGGCTGCCAAC<br>*HaA.R2 TAACGGTACTTGGCAGTMAG<br>Haa.R2.ducks CAGCCGCCACCTTCTTGCC<br>Haa.F2.ducks CCCAGACCAAGACCTACTTCCCC<br>*LCAT.2F GTGGTGAACTGGATGTGCTACCG<br>*LCAT.3F GTACCTGGCTTYGGCAAGACC<br>LCAT.3FB GGTGCAYATCCGYGTNCCTGG<br>*LCAT.3R ACCTGCCAGTTTGCTCTGGTCCAG<br>LCAT.4F AAYGGCTAYGTGAGGGACCA<br>LCAT.4R CCRACYCTCCAGTCATARGG<br>*LCAT.5R GCACCCAGNGAGATGAAGCC<br>LCAT.5Rb CCCGATGTACTGATCTTTCCAGG<br>PEPCK3.Fi TGCAGCAGATAGCAARTGAGGTG<br>PEPCK3.Ri CTGYAGTAAAGGTGGGTGGAGG<br>*PEPCK3F GGTCGCTGGATGTCAGAAGAGG<br>PEPCK3F.2 TCAATACCAGATTCCCAGGCTGC<br>*PEPCK3R CCATGCTGAAGGGGATGACATAC<br>GTP1601F ACGAGGCCTTTAACTGGCAGCA<br>GTP1793R CTTGGCTGTCTTTCCGGAACC<br>*PEPCK9F GGAGCAGCCATGAGATCTGAAGC<br>PEPCK9F.2 CTTACATTTTCTGTTCTGCTAGAGC<br>*PEPCK9R GTGCCATGCTAAGCCAGTGGG<br>PEPCK9R.2 CTTGAGAGCTGGCTTTCATTG | *L1263 YAAAGCATGRCACTGAA<br>L1267 YAAAGCATGRCACTGAAGHYG<br>L1512 TAAGCAATGAGTGHAARCTYGACTTAG<br>L1753 AAACTGGGATTAGATACCCCACTAT<br>*L1754 TGGGATTAGATACCCCACTATG<br>L1843 AAACYCTAAGGACYTGGCGG<br>L1936 CAGCCTAYATACCGCCGTC<br>L2010 TARHAMGACAGGTCRAGGTATAGC<br>H1530 GTGGCTGGCACARGATTTACC<br>H1806 GTTTYAAGCGTTKGYGCTCGTA<br>H1858 TCGATTATAGAACAGGCTCCTCTAG<br>*H1859 TCGDTTRYAGRACAGGCTCCTCTA<br>H1918 GACGGCGGTATRTAGGCTG<br>H1993 DDGCTATACCTYGACCTGTC<br>H2084 NTTTACTDCTAAATCCDCCTT<br>*H2294 TYTCAGGYGTARGCTGARTGCTT |

**References:**

Bock, W. J. History and nomenclature of avian family-group names. *Bulletin of the American Museum of Natural History* **222**, 1–281 (1994).

Buckner, J. C., Ellingson, R., Gold, D. A., Jones, T. L. & Jacobs, D. K. Mitogenomics supports an unexpected taxonomic relationship for the extinct diving duck *Chendytes lawi* and definitively places the extinct Labrador Duck. *Mol. Phylogenetics Evol.* **122**, 102–109 (2018).

Bulgarella, M., Sorenson, M. D., Peters, J. L., Wilson, R. E. & McCracken, K. G. Phylogenetic relationships of *Amazonetta*, *Speculanas*, *Lophonetta*, and *Tachyeres*: four morphologically divergent duck genera endemic to South America. *J. Avian Biol.* **41**, 186–199 (2010).

*Clements, J. F., P. C. Rasmussen, T. S. Schulenberg, M. J. Iliff, T. A. Fredericks, J. A. Gerbracht, D. Lepage, A. Spencer, S. M. Billerman, B. L. Sullivan, and C. L. Wood. The eBird/Clements checklist of Birds of the World: v2023 (2023).*

Donne-Goussé, C., Laudet, V. & Hänni, C. A molecular phylogeny of anseriformes based on mitochondrial DNA analysis. *Mol. Phylogenetics Evol.* **23**, 339–356 (2002).

García-Moreno, J., Sorenson, M. D. & Mindell, D. P. Congruent avian phylogenies inferred from mitochondrial and nuclear DNA sequences. *J. Mol. Evol.* **57**, 27–37 (2003).

Gonzalez, J., Düttmann, H. & Wink, M. Phylogenetic relationships based on two mitochondrial genes and hybridization patterns in Anatidae. *J. Zoöl.* **279**, 310–318 (2009).

Johnson, K. P. & Sorenson, M. D. Phylogeny and Biogeography of Dabbling Ducks (Genus: Anas): A Comparison of Molecular and Morphological Evidence. *Auk* **116**, 792–805 (1999).

Livezey, B. C. A phylogenetic classification of waterfowl (Aves: Anseriformes), including selected fossil species. *Annals of Carnegie Museum* **66**, 457–496 (1997).

McCracken, K. G. & Sorenson, M. D. Is homoplasy or lineage sorting the source of incongruent mtDNA and nuclear gene trees in the stiff-tailed ducks (*Nomonyx-Oxyura*)? *Syst. Biol.* **54**, 35–55 (2005).

McCracken, K. G. *et al.* Parallel evolution in the major haemoglobin genes of eight species of Andean waterfowl. *Mol. Ecol.* **18**, 3992–4005 (2009).

Mindell, D. P. *et al.* Phylogenetic relationships among and within select avian orders based on mitochondrial DNA. in *Avian Molecular Evolution and Systematics* (ed. Mindell, D. P.) 213–247 (1997).

Sorenson, M. D., Oneal, E., García-Moreno, J. & Mindell, D. P. More taxa, more characters: the hoatzin problem is still unresolved. *Mol. Biol. Evol.* **20**, 1484–1498 (2003).

Sun, Z. *et al.* Rapid and recent diversification patterns in Anseriformes birds: Inferred from molecular phylogeny and diversification analyses. *PLoS ONE* **12**, e0184529 (2017).

*Swofford, D. L. PAUP*. Phylogenetic Analysis Using Parsimony (*and Other Methods). Version 4. Sinauer Associates, Sunderland, Massachusetts (2002).*

Winkler, D. W., Billerman S. M., Lovette, I. *Bird Families of the World: An Invitation to the Spectacular Diversity of Birds*. Lynx Edicions (2015).

### Appendix IV: Waterfowl Morphometrics and Trait Evolution

#### Characterization of Skull and Hindlimb Shape

PC of Skull. – PC1 for skull shape (35.3%) describes a suite of proportions of the skull and rostrum (Fig. 3). Most notably the length and curvature of the rostrum anterior to the nares, with low values indicating a relatively short bill and high values indicating a relatively long bill. The width and height of the skull are also associated with PC1, low values have a wider skull (particularly across the posterior base of the bill), and a taller more domed skull. Higher values having a much narrower and low, sloped skull. The extreme lower values of PC1 are represented by Anhimidae which are highly aberrant compared to other anseriforms. The next lowest values are represented by species normally referred to as "geese," including several distantly related lineages (i.e., members of *Anser, Branta, Chloephaga, Cereopsis, Nettapus,* etc.). All straining/surface and piscivorous groups have relatively high PC1 values but the highest values are represented by "shovelers" (*Spatula*) with extremely long and wide bills.

PC2 for skull shape (19.4%) largely describes the dorsoventral flexion of the skull, characterised by the angle between the top of the supraoccipital and the tip of the rostrum (Fig. 3). Species with low PC2 scores have a ventrally flexed skull when compared to the position of the supraoccipital. Species with higher PC2 values have little to no flexion of the skull. PC2 also reflects the width of the skull as measured across the quadrates, with high values representing wider skulls and lower values representing narrower skulls. Species with low PC2 also have relatively smaller nares compared to species with higher PC2 values. The mergansers (*Mergus Lophodytes, Mergellus*) have among the highest PC2 values, having particularly horizontally orientated skulls with wide set quadrates. *Biziura lobata* also has a notably high PC2 value. *Tadorna tadorna* has a notably high PC2 value compared to other surface straining species, but this might be due to the highly upturned bill which would reduce the angle between the supraoccipital and tip of the bill. Lower PC2 values are not represented by a singular ecomorphotype, the lowest values are observed in species such as *Nettapus pulchellus, Malacorhynchus membranaceus,* and *Chloephaga melanoptera*.

PC of Femur. – PC1 for femur shape (30.8%) largely describes the curvature and size of the crista trochanteris (Fig. 4A). Low PC1 values have a more strongly curved femur from the midpoint of the shaft and a proximal end with a low profile, making the crista trochanteris less prominent. High PC1 values have a straighter femoral shaft, with the femoral condyles rising above the shaft in lateral view. The crista trochanteris is also deeper and more prominent. Low PC1 values are largely representative of diving species, and this curvature has been noted as a shared characteristic of divers previously (McCracken 1999). Lower scores are largely terrestrial grazing species.

PC2 (24.2%) is characterised by the width of the distal shaft, the position of the tubercle on the posterior face of the shaft, the length and orientation of the tuberculum m. gastrocnemialis lateralis (tgl), the extent of curvature at the very distal end portion of the femur, and the length of the articular surfaces of the condyles (Fig. 4A). High PC1 scores have a mediolaterally narrow distal shaft, a more distally located posterior tubercle, a longer and more vertically orientated tgl, a more severely curved distal portion, and a longer articular surface. Lower scores have a more proximally placed posterior tubercle, a shorter and more horizontally orientated tgl, a straighter femur, and shorter articular surfaces. The highest PC1 scores are almost entirely occupied by diving-grasping species such as *Tachyeres pteneres* and *Melanitta fusca*. The lowest scores are occupied by Anhimidae and *Anseranas semipalmata*.

PC of Tibiotarsus. – PC1 for tibiotarsus shape (41.5%) describes the length of the shaft, cnemial crest, and fibular crest as well as the width of the condyle (Fig. 4B). High PC1 scores indicate a shorter shaft, taller cnemial crest, longer fibular crest, and a wider condyle. Low PC1 scores indicate a longer shaft, shorter cnemial crest, shorter fibular crest, and a narrower condyle. Diving species tend to have higher PC1 scores while there is little differentiation between other groups.

PC2 (14.02%) describes the width of the proximal articular face and the width of the vertical blade of the cnemial crest (Fig 4B). Higher scores have a wider articular face and wider cnemial crest, and lower scores have a narrower articular face with a narrow cnemial crest. Diving species largely occupy the higher scores, but other groups also heavily overlap with these scores. Diving species are the only group that occupy a distinct place in the PC1-PC2 morphospace, having generally shorter, wider tibiotarsi with prominent cnemial crests.

PC of Tarsometatarsus. – PC1 for tarsometatarsus shape (55.7%) largely describes the relative length of the shaft as well as the relative width and position of the II trochlea (Fig 4C). High PC1 scores represent species with shorter and stouter tarsometatarsi with a proximally placed II trochlea. Low PC1 scores represent species with longer and thinner tarsometatarsi, with more distally placed II trochlea. High PC1 scores are observed in diving species, with *Biziura lobata, Oxyura* spp. and *Tachyeres pteneres* having the highest scores. Wading species have by far the lowest PC1 scores, with terrestrial grazers having the next lowest scores.

PC2 (9.2%) describes the size of the crista medialis hypotarsi (cmh) and angle of the trochlea (Fig 4C). Higher PC2 scores have a longer cmh with the distal ends of the trochlea facing more distally. Lower PC2 scores have a shorter cmh with the distal ends of the trochlea facing more medially. The various ecological categories are spread across PC2, though some weak correlation is evident. Diving species generally have higher PC2 scores and terrestrial species generally lower scores but there is significant overlap. *Anas chlorotis* has a notably high PC2 score, and *Merganetta armata* has a less extreme but still notably high PC2 score.

### Appendix V: Phenotype Transitions and Rates of Phenotypic Evolution

**Supplementary Figure 6.** *a priori* Ornstein-Uhlenbeck models for the skull (A), femur (B), tibiotarsus (C), and tarsometatarsus (D). Black circles represent significant transitions in phenotype. Branches are colored according to each transition.

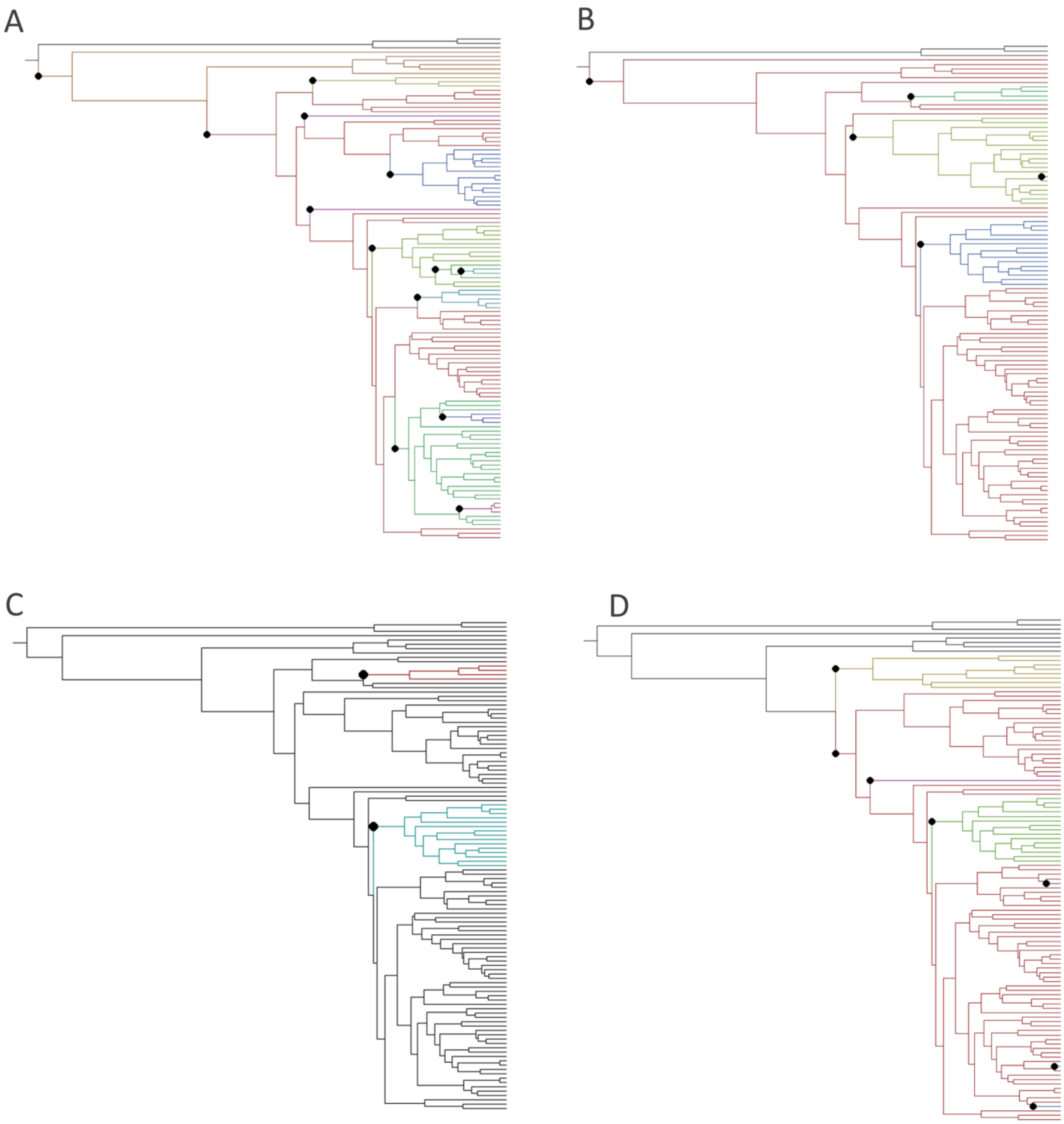

Appendices

**Supplemental Figure 7.** Rates of phenotypic evolution across a time-calibrated waterfowl phylogeny. Evolutionary rates of the skull (A), tibiotarsus (B), femur (C), tarsometatarsus (D). Black circles represent a significant shift toward increasing evolutionary rates. Red circles represent a significant shift toward decreasing evolutionary rates. All elements shown are 3D meshes of *Hymenolaimus malacorhynchus (*NMNH 19024).

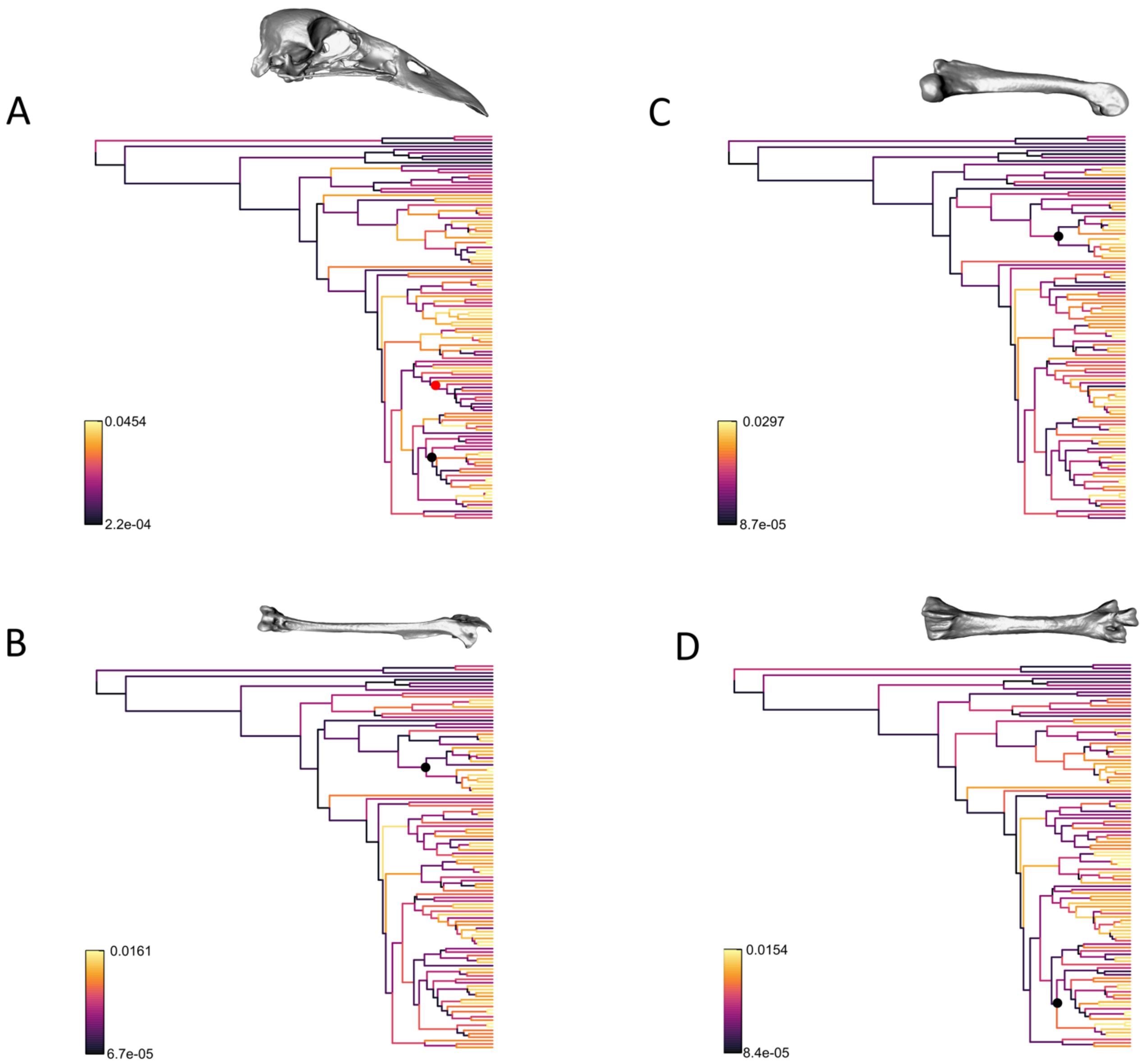

**Appendix VI: Morphological convergence within diving invertivores and surface strainers**

We found strong support for convergent evolution among the large macroinvertivore species, such as *Somateria molissima, Tachyeres pteneres*, and *Biziura lobata*. This group has been proposed before (McCracken et al. 1999), however we also found them convergent in skull shape with the riverine species *Hymenolaimus malacorhynchus* and *Merganetta armata*. The features of their skull shape which unite them, a deeper and taller bill, and wider posterior skull are all associated with applying a more efficient bite (Herrel et al. 2008). They also share a more horizontally orientated skull with forward-facing eyes needed for prey detection and capture (Martin et al. 2007; Cantalay et al. 2023). There are also morphological distinctions in femur morphology from divers which strain from freshwater versus those capturing moving or larger prey. Though foraging alone was the strongest factor in femur shape evolution there were still notable differences across diet. For example, the size of the tuberculum m. gastrocnemialis lateralis was consistently larger in diving grasping species than the diving strainers such as *Oxyura jamaicensis*. The tuberculum m. gastrocnemialis lateralis is associated with the gastrocnemius muscle which helps drive the power stroke in diving (Clifton et al. 2018). This muscle is very large in many diving birds such as loons, grebes and cormorants (Clifton et al. 2018). Perhaps the coastal environments and fast flowing rivers require a more powerful stroke to control direction in more turbulent waters, and for coastal species to be able to dive to deeper depths.

While united within this study, the diving invertivore species seem to represent two distinct ecological groups, the large bodied macroinvertivores and the smaller mountainous river species which feed on smaller invertebrates. The ecological similarity between *H. malacorhynchus* and *M. armata* have been remarked upon before along with the other mountainous species *Salvadorina waiguensis* (Kear 1975; Veltman 1990). Along PC1 and PC2 for skull shape, the axes most clearly associated with diet, *S. waiguensis* plotted within the variation of the diving graspers suggesting that this species might be converging upon this ecotype as well.

We also found the surface feeding strainers, often called dabblers, to be convergent in skull shape and femur shape. This contrasts with hypotheses from the paleontological literature (Worthy and Lee 2008; Zelenkov 2020), which suggests that dabbling is the ancestral condition of anatids, and thus homologous. These results might be due to the lack of fossil species in our study strongly influencing the ancestral state reconstruction in our morphospace. Brownian motion models, as is used in ancestral state reconstruction in conevol 'averages' shape between descending tips (Stayton 2015). This form of ancestral state reconstruction can potentially be inaccurate (Kuo et al. 2023) and highlights the need to include fossil taxa. While some dabbling species may be convergent in morphology, such as the spatulate billed *Malacorhyncus membranaceus* and *Spatula* we find it unlikely that the entire ecotype is convergent, and instead represents a natural variation of a similar ancestral condition.

Appendices

### References


Cantlay, J. C., Martin, G. R., McClelland, S. C., Potier, S., O'Brien, M. F., Fernández-Juricic, E., Bond, A. L., & Portugal, S. J. (2023). Binocular vision and foraging in ducks, geese and swans (Anatidae). *Proceedings of the Royal Society B: Biological Sciences*, *290*(2006), 20231213. https://doi.org/10.1098/rspb.2023.1213

Clifton, G. T., Carr, J. A., & Biewener, A. A. (2018). Comparative hindlimb myology of foot-propelled swimming birds. *Journal of Anatomy*, *232*(1), 105–123. https://doi.org/10.1111/joa.12710

Herrel, A., Podos, J., Huber, S. K., & Hendry, A. P. (2005). Evolution of bite force in Darwin's finches: A key role for head width. *Journal of Evolutionary Biology*, *18*(3), 669–675. https://doi.org/10.1111/j.1420-9101.2004.00857.x

Kear, J. (1975). Salvadori's Duck of New Guinea. *Wildfowl*, *26*(26), Article 26.

Kuo, P.-C., Benson, R. B. J., & Field, D. J. (2023). The influence of fossils in macroevolutionary analyses of 3D geometric morphometric data: A case study of galloanseran quadrates. *Journal of Morphology*, *284*(6), e21594. https://doi.org/10.1002/jmor.21594

Martin, G. R., Jarrett, N., & Williams, M. (2007). Visual fields in Blue Ducks Hymenolaimus malacorhynchos and Pink-eared Ducks Malacorhynchus membranaceus: Visual and tactile foraging. *Ibis*, *149*(1), 112–120. https://doi.org/10.1111/j.1474-919X.2006.00611.x

McCracken, K. G., Harshman, J., McClellan, D. A., & Afton, A. D. (1999). Data Set Incongruence and Correlated Character Evolution: An Example of Functional Convergence in the Hind-Limbs of Stifftail Diving Ducks. *Systematic Biology*, *48*(4), 683–714. https://doi.org/10.1080/106351599259979

Stayton, C. T. (2015). The definition, recognition, and interpretation of convergent evolution, and two new measures for quantifying and assessing the significance of convergence. *Evolution*, *69*(8), 2140–2153. https://doi.org/10.1111/evo.12729

Veltman, C. J., Collier, K. J., Henderson, I. M., & Newton, L. (1995). Foraging ecology of blue ducks *Hymenolaimus malacorhynchos* on a New Zealand river: Implications for conservation. *Biological Conservation*, *74*(3), 187–194. https://doi.org/10.1016/0006-3207(95)00029-4

Worthy, T. H., & Lee, M. S. Y. (2008). Affinities of Miocene Waterfowl (Anatidae: Manuherikia, Dunstanetta and Miotadorna) from the St Bathans Fauna, New Zealand. *Palaeontology*, *51*(3), 677–708. https://doi.org/10.1111/j.1475-4983.2008.00778.x

Zelenkov, N. V. (2020). Cenozoic Evolution of Eurasian Anatids (Aves: Anatidae s. l.). *Biology Bulletin Reviews*, *10*(5), 417–426. https://doi.org/10.1134/S2079086420050096
